# Extensive SARS-CoV-2 spike sequence diversification converges on a bounded, low-dimensional physicochemical space

**DOI:** 10.64898/2026.11.12.766201

**Authors:** Yiannis Manoussopoulos, Cleo Anastassopoulou, Marianna Manoussopoulou, Constantinos Siettos, Julian A. Hiscox, John P.A. Ioannidis, Athanasios Tsakris

**Affiliations:** Department of Microbiology, Medical School, National and Kapodistrian University of Athens, Athens, Greece; ELGO-Demeter, Plant Protection Division of Patras, Laboratory of Virology, Patras, Greece; Dipartimento di Matematica e Applicazioni ‘Renato Caccioppoli’, Università degli Studi di Napoli Federico II, Naples, Italy; Department of Infection Biology and Microbiomes, Institute of Infection, Veterinary and Ecological Sciences, University of Liverpool, Liverpool, United Kingdom; Departments of Medicine, Epidemiology and Population Health, and Biomedical Data Science, Stanford University, Stanford, CA, United States of America

**Keywords:** SARS-CoV-2, spike protein, phenotypic clustering, physicochemical properties, Shannon entropy, viral evolution

## Abstract

During the COVID-19 pandemic, SARS-CoV-2 accumulated extensive amino acid variation in its spike protein. However, the extent to which sequence diversification translates into physicochemical diversity remains unclear. Here, we analyzed 271,769 unique spike protein sequences from 12 geographical regions using three physicochemical descriptors: net charge at pH 7.0, hydropathy, and aliphatic index. To characterize temporal changes in physicochemical space, we compared the squared Mahalanobis distances of observed sequences with those of Monte Carlo-simulated sequences containing randomized amino acid substitutions, constrained to the number of empirically variable positions. We applied trimmed clustering to characterize sequence distributions in physicochemical space and used Shannon entropy to assess spatiotemporal relationships between genotypic and physicochemical diversity across geographic regions. Empirical physicochemical displacement increased early in the observation period and subsequently stabilized, despite continued increases in variable amino acid positions and accumulated substitutions. In contrast, simulated trajectories exhibited greater and sustained physicochemical divergence. Trimmed clustering identified six distinct physicochemical groups repeatedly occupied by genetically divergent lineages differing in mutation burden, amino acid composition, and emergence time. Shannon entropy analyses revealed broadly similar nonlinear temporal patterns across geographic regions, with differences between genotypic and physicochemical entropy tending to be negative. These findings indicate that extensive spike protein sequence diversification occurred within a comparatively restricted physicochemical space, consistent with a many-to-one relationship between amino acid sequences and physicochemical states. This framework provides a quantitative approach for investigating physicochemical constraints on viral protein evolution and may inform future machine learning approaches to epitope prediction and vaccine antigen design.

## Introduction

The severe acute respiratory syndrome coronavirus 2 (SARS-CoV-2) spike protein is the principal surface glycoprotein mediating host-cell recognition and viral entry (1, 2). During viral evolution, the spike protein accumulates amino acid substitutions under multiple selective pressures, particularly those associated with angiotensin-converting enzyme 2 (ACE2) receptor binding (3–5) and antibody-mediated immune escape (6, 7). These substitutions can alter receptor-binding affinity and protein structure, thereby modulating viral entry, immune evasion, and overall viral fitness (5, 6).

Throughout the COVID-19 pandemic, the SARS-CoV-2 spike protein evolved from relatively limited sequence variation in early 2020 to highly divergent lineages by 2025 relative to the Wuhan-Hu-1 reference sequence. Among the earliest changes associated with increased viral fitness and transmission were the D614G substitution in spike and the linked P323L substitution in the RNA-dependent RNA polymerase (NSP12) (8–10). SARS-CoV-2 variants have been classified using several complementary nomenclature systems, including the Pango lineage system. Variants carrying mutations associated with, or predicted to affect, transmissibility, disease severity, or immune evasion that were likely to have a population-level impact were designated as variants of concern (VOCs). These included Alpha, Delta, and Omicron, whose descendants subsequently diversified into numerous lineages that circulated globally.

This diversification reflects selection acting on combinations of mutations that influence immune evasion, receptor engagement, and viral fitness (6). Rather than acting independently, amino acid substitutions accumulate in constellations whose combined effects can alter the physicochemical properties of the spike protein (3, 11, 12). Consequently, phenotypic characteristics, including receptor binding, immune escape, and viral adaptation, may depend not only on individual substitutions but also on their collective effects on the protein’s physicochemical landscape. Previous studies have investigated specific physicochemical properties of spike, including variation in electrostatic charge among SARS-CoV-2 lineages (13, 14). In addition, unsupervised deep learning approaches have demonstrated that sequence-based representations can reveal population structure among SARS-CoV-2 strains (15). However, the relationship between extensive sequence variation and multidimensional physicochemical diversity remains insufficiently characterized.

Specifically, it remains unclear whether extensive spike sequence diversification has generated a correspondingly broad physicochemical landscape or whether genetically divergent lineages occupy a more restricted repertoire of physicochemical states. Resolving this question may clarify how sequence variation translates into physicochemical variation and how the relationship between genotypic and physicochemical diversity changes across geographic regions and over time.

To address this gap, we integrated Monte Carlo simulations of memory-dependent and memoryless physicochemical divergence, multidimensional clustering of spike protein physicochemical properties, and Shannon entropy analyses to characterize temporal relationships between genotypic and physicochemical diversity across multiple geographic regions. Our analyses indicate that, extensive spike sequence diversification occurred within a comparatively restricted multidimensional physicochemical space.

Genetically divergent lineages repeatedly occupied distinct physicochemical groups, while overall physicochemical displacement stabilized despite continued accumulation of amino acid variation. These findings support a many-to-one relationship between spike sequence variation and physicochemical states and provide a quantitative framework for investigating potential physicochemical constraints on viral protein evolution.

## Materials and Methods

### Software

Dataset preparation, physicochemical analyses, statistical analyses and data visualization were performed in R (16). Analyses utilized functions from base R and the following packages: Biostrings (17), boot (18) cluster (19), data.table (20), FactoMineR (21), factoextra (22), ggplot2 (23), glmnet (24), Peptides (25), tidyverse (26), tidyr (27), tclust (28), mgcv (29), openxlsx (30), and plotly (31). Multiple sequence alignments were generated using the MAFFT algorithm (32) through the R interface.

### Data retrieval, cleaning, and sequence alignment

Complete SARS-CoV-2 protein sequence records were retrieved from the Global Initiative on Sharing All Influenza Data (GISAID) database on January 16, 2026, and records encoding the spike (S) protein were extracted (33). Sequences were processed and validated using a multistep pipeline (Supplementary Table 1). Records containing ambiguous amino acid residues or incomplete metadata in their FASTA headers were excluded. The remaining records were further screened to ensure consistency across metadata fields.

Processed sequences were grouped by country, collection date, and sequence identity, and the corresponding frequencies were calculated. Within each country, records were ordered chronologically by collection month and year, and the frequencies of duplicate S protein sequence were determined before deduplication. Identical sequences reported from different countries were retained. Metadata absent from the FASTA headers were manually retrieved using the corresponding GISAID accession identifiers and incorporated into the dataset. Records with collection dates preceding the first reported appearance of their assigned SARS-CoV-2 variant were excluded.

For each country, curated unique sequences were exported in FASTA format and aligned using the MAFFT algorithm, with the GISAID (earliest) sequence hCoV-19/Wuhan/WIV04/2019 (WIV04; EPI_ISL_402124) used as the reference. The resulting alignments were converted to comma-separated values (CSV) format, and amino acid positions were renumbered according to the reference sequence to maintain consistency with the original residue numbering.

Physicochemical properties were calculated for each aligned S protein sequence, using the *Peptides* R package. Net charge at pH 7.0 was calculated using the EMBO charge scale (34, 35), hydropathy using the Kyte-Doolittle index (36), and the aliphatic index as a measure of thermostability(37). The Boman index and instability index were also calculated as indicators of protein binding potential (38) and protein stability (39), respectively. Pairwise associations among physicochemical properties were assessed using Spearman’s rank correlation. Net charge at pH 7.0, hydropathy, and aliphatic index were treated as complementary dimensions of a multidimensional physicochemical phenotype, with each S protein sequence represented as a point in physicochemical space.

Twelve geographic regions were included in the analyses: ten countries (Australia, Brazil, Canada, France, Germany, India, Italy, Japan, South Africa, and the United Kingdom) and two U.S. states (California and New York). Regions were selected primarily on the basis of the availability of large sequence collections with sustained temporal coverage, supplemented by locations associated with the emergence of major SARS-CoV-2 variants of concern.

Pango-designated lineages were grouped according to the World Health Organization (WHO) Greek-letter classification of SARS-CoV-2 variants. Omicron lineages were further subdivided into ten groups representing distinct evolutionary and biological categories (Supplementary Table 2). This grouping was informed by published evidence supporting the distinct evolutionary and immunological characteristics of BA.1 (40), BA.2 (40), BA.4/BA.5 (41, 42), XBB (43), and BA.2.86/JN.1 (44) represent distinct evolutionary and immunologic groups (40–44). Lineages predating the introduction of the WHO Greek-letter nomenclature were classified as Unassigned.

### Displacement analysis

To quantify the displacement of S protein physicochemical properties associated with natural selection relative to expectations under conditional randomization, three Monte Carlo null models were implemented (45). In the first, a memoryless model, the empirically observed variable positions of each sequence were retained, while amino acid identities at these positions were independently randomized across successive year-months. In the second, a memory-based model, the empirically observed variable positions were likewise retained, but randomized amino acid outcomes at recurrent variable positions were propagated across successive year-months. In the third, a whole-spike memoryless model, only the number of distinct variable positions in each empirical sequence was retained, while their locations were randomized across the entire S protein sequence. In all three models, amino acids were sampled uniformly from the 20 standard amino acids (p=1/20), thereby avoiding the introduction of additional empirical constraints beyond those explicitly imposed by each randomization scheme. For each year-month, up to 300 records were randomly sampled without replacement, and 200 Monte Carlo realizations were generated across the complete time series.

Displacement from the baseline physicochemical phenotype was quantified using the squared Mahalanobis distance (*d*^2^) (46). Distances were calculated for observed and simulated sequences in two complementary ways: (i) by year-month to characterize temporal displacement and (ii) within fixed-width bins of empirically observed amino acid changes independently of calendar time, to characterize displacement as a function of sequence divergence.

For the latter analysis, simulated sequences were grouped according to the number of amino acid changes on their corresponding empirical sequences.

The baseline physicochemical phenotype was defined using S protein sequences collected during the first three months of the pandemic (n=815). For each sequence *i, d*^2^ was calculated as:

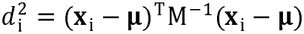

where ***x***_i_, the physicochemical feature vector for sequence i, comprising net charge at pH 7.0, hydropathy, and aliphatic index; **μ** is the mean feature vector of the baseline population; and M is the corresponding covariance matrix. The squared Mahalanobis distance quantifies multivariate displacement from the baseline phenotype while accounting for covariance among physicochemical properties.

### PCA clustering and validation

Principal component analysis (PCA) was used as an exploratory method to characterize variance and identify the principal axes of variation in physicochemical space (47, 48), using net charge at pH 7.0, hydropathy, and aliphatic index as input variables. Prior to analysis, all variables were centered and scaled to zero mean and unit variance using *scale* function in R. PCA was subsequently performed on the standardized variables using the *prcomp* function.

Robust clustering was performed using the *tclust* package to identify recurrent physicochemical phenotypes independently of viral lineage assignments. This method accommodates potential outliers through a trimming parameter (**α**) and constrains the eigenvalue ratios of cluster covariance matrices using a restriction factor, thereby controlling cluster geometry and limiting excessive heterogeneity in cluster dispersion (28, 49, 50).

The *ctlcurves* function in the *tclust* package was used to evaluate candidate trimming levels (**α**) and numbers of clusters (k) based on classification trimmed likelihood (CTL) curves, incremental improvement in CTL (ΔCTL), and clustering stability, following established approaches to robust clustering and model validation (28, 49, 51, 52) . Seven trimming levels (α = 0, 0.002, 0.005, 0.01, 0.02, 0.03, and 0.05) and 25 candidate cluster numbers (k = 1–25) were evaluated. For each parameter combination, four independent batches of 30,000 randomly selected sequences were analyzed, with sampling stratified by collection year-month and WHO variant designation. The incremental improvement associated with increasing *k* was calculated as the change in CTL between successive models (ΔCTL). Clustering stability across subsamples was assessed using the adjusted Rand index (ARI) (53).

The final number of clusters was selected by jointly considering model fit, clustering stability, physicochemical resolution, and temporal interpretability (54). Particular attention was given to candidate solutions containing five to seven clusters (k = 5-7).

Following selection of **α** and k, the covariance matrix restriction factor was optimized separately through sensitivity analysis. Candidate restriction factors were compared in terms of CTL and clustering stability across subsamples, and the selected value was used to constrain the covariance matrices of the final clustering model (28, 49).

To assess the robustness of cluster assignments to potential geographic sampling bias, ridge regression was performed using the *glmnet* package (α = 0) to evaluate whether variation in sequencing intensity across countries was associated with cluster assignment (24). Associations between cluster assignments and geographic regions were quantified using Cramér’s V statistic (55)

### Cluster statistics

Within-cluster variability in physicochemical properties was characterized using non-parametric bootstrap resampling implemented with the *boot* package in R (56). For each cluster, 1,000 bootstrap replicates were generated by sampling sequences with replacement from the original cluster data. For each replicate, the mean, standard deviation (SD), median, and interquartile range (IQR) were calculated for net charge at pH 7.0, hydropathy, and aliphatic index. The resulting bootstrap distributions were used to estimate 95% confidence intervals (CIs) for each summary statistic.

Cluster separation and robustness were further evaluated using multivariate analysis of variance (MANOVA), pairwise comparisons of cluster centroids, and linear discriminant analysis (LDA) (57). To assess centroid stability, 200 bootstrap resamples were generated with replacement, and cluster centroids were recalculated using the three standardized physicochemical variables.

The effect of within-country sequence deduplication on clustering results was assessed by comparing deduplicated and frequency-weighted datasets. Differences between the resulting solutions were evaluated by quantifying shifts in cluster centroids and comparing the proportion of variance explained by PCA in the weighted and unweighted datasets.

### Shannon entropy of variants and clusters

Shannon entropy (58) was used to quantify the temporal diversity of genotypic variants and phenotypic clusters at monthly intervals, enabling direct comparison of genotypic and phenotypic diversity over time. For each geographic region (*r*) and month (*t*), the total number of sequences was calculated as 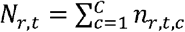 where *n*_r,t,c_ is the number of sequences assigned to cluster or variant *c*, and *C* is the total number of genotypic variant or phenotypic cluster categories.

The proportion of sequences belonging to each cluster or variant was calculated as 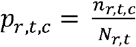 . Shannon entropy was then computed for each region-month combination as:

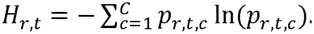

Temporal entropy trajectories were visualized using locally estimated scatterplot smoothing (LOESS) implemented in the *ggplot2* package. To reduce potential bias associated with unequal monthly sample sizes, entropy was additionally estimated using equal-sample-size resampling. Region-month combinations containing at least 30 sequences were eligible for resampling. For each eligible region-month, B= 1,000 independent random subsamples of N=30 sequences were drawn without replacement. Region-month combinations containing fewer than 30 sequences were not resampled and represented as gaps or dotted lines in the temporal figures.

Genotype–phenotype decoupling was quantified as the difference between phenotypic and genotypic Shannon entropy: 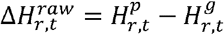, where superscripts *p*, and *g* denote phenotypic (cluster) and genotypic (variant) diversity, respectively. To account for differences in the numbers of available categories, entropy was normalized by the logarithm of the number of observed categories (59).

For each resampling replicate (b), normalized phenotypic and genotypic entropy were calculated as: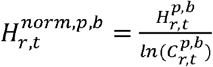, for phenotype and 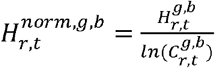 for genotype, where 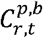, and 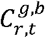 denote the numbers of phenotypic cluster categories and genotypic variant categories observed in resample b, respectively. When only one category was represented in a subsample, normalized entropy was defined as zero.

Normalized genotype–phenotype decoupling for each subsample was calculated as 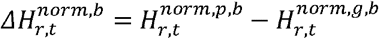 The final normalized genotype–phenotype entropy difference for each region-month combination was obtained by averaging across the B resamples: 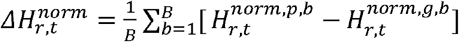. Positive values of 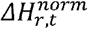 indicate greater normalized phenotypic than genotypic diversity, whereas negative values indicate greater normalized genotypic diversity.

Temporal variation in the magnitude of genotype-phenotype decoupling was modelled using generalized additive models (GAMs) implemented in the *mgcv* package. The absolute normalized entropy difference |*ΔH*| was used as the response variable, with time included as a smooth predictor and geographic region as a parametric effect. The model was specified as:

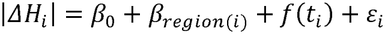

where | Δ*H*| is the response variable representing the magnitude of genotype-phenotype decoupling of observation *i, β*_0_ is the intercept, *β*_region(i)_ is the parametric effect associated with geographic region of observation *i, f*(*t*_i_) is a smooth function of time, *t*_i_ denotes the observation time, and *ε*_i_ is the residual error. To account for temporal dependence between successive monthly observations, an autoregressive correlation structure of order 1 [AR(1)] was applied to residuals within each geographic region. Smoothing parameters were estimated using restricted maximum likelihood (REML).

## Results

### Isolation of unique sequences for analysis

A total of 17,113,342 SARS-CoV-2 spike protein sequences were retrieved from GISAID and subjected to quality control (Supplementary Table 1). Of these, 6,815,427 records were excluded because of ambiguous amino acid entries, non-human host origin, insufficient sequences length, or ambiguous country annotations, leaving 10,297,915 records for subsequent analysis. Within-country deduplication further removed 9,632,887 duplicate sequences, resulting in 665,028 unique records. From this dataset, 272,985 sequences originating from the 12 regions included in this study were retained for downstream analyses (Table 1). The annual distribution of unique spike protein sequences included for each country and region is presented in Supplementary Table 3. Finally, sequences with missing variant annotations or inconsistent collection dates were excluded, and the resulting country- and region-specific datasets were prepared for the corresponding downstream analyses (Supplementary Table 4).

**Table 1.** SARS-CoV-2 spike sequences included in the study, by country or U.S. state, before and after sequential filtering steps. Identical amino acid sequences were deduplicated within each geographical region and year-month. Sequences with missing variant assignments, as determined by distinct amino acid combinations, were subsequently excluded, followed by exclusion of sequences with missing variant labels. Percentages were calculated relative to the number of sequences before deduplication.

|  | DEDUPLICATION |  |  |  |  |
|---|---|---|---|---|---|
| COUNTRY/REGION | BEFORE (n) | AFTER (n, %) | DUPLICATES (n, %) | MISSING (n) | RETAINED* (n) |
| Australia | 135,980 (100.0) | 10,520 (7.7) | 125,460 (92.3) | 8 | 10,512 |
| Brazil | 152,969 (100.0) | 16,936 (11.1) | 136,033 (88.9) | 180 | 16,756 |
| Canada | 604,583 (100.0) | 30,440 (5.0) | 574,143 (95.0) | 388 | 30,052 |
| France | 384,122 (100.0) | 31,816 (8.3) | 352,306 (91.7) | 126 | 31,690 |
| Germany | 447,167 (100.0) | 33,291 (7.4) | 413,876 (92.6) | 71 | 33,220 |
| India | 68,566 (100.0) | 13,716 (20.0) | 54,850 (80.0) | 27 | 13,689 |
| Italy | 99,247 (100.0) | 12,938 (13.0) | 86,309 (87.0) | 58 | 12,880 |
| Japan | 622,179 (100.0) | 26,580 (4.3) | 595,599 (95.7) | 115 | 26,465 |
| South Africa | 16,161 (100.0) | 2,930 (18.1) | 13,231 (81.9) | 24 | 2,906 |
| USA (California) | 493,453 (100.0) | 23,890 (4.8) | 469,563 (95.2) | 60 | 23,830 |
| USA (New York) | 265,042 (100.0) | 14,964 (5.6) | 250,078 (94.4) | 77 | 14,887 |
| United Kingdom | 2,008,228 (100.0) | 54,964 (2.7) | 1,953,264 (97.3) | 82 | 54,882 |
| **Total** | **5,297,697** | **272,985** | **5,024,712** | **1,216** | **271,769** |

\* Retained denotes deduplicated sequences remaining after exclusion of sequences with missing variant labels.

### Selection of physicochemical variables

Pairwise correlation analysis revealed a strong negative correlation between the Boman index and hydropathy (ρ=-0.82), whereas the aliphatic index and charge exhibited a moderately strong positive correlation (ρ=0.61). In contrast, weak correlations were observed between charge and hydropathy (ρ=-0.13), charge and aliphatic index (ρ=-0.16), and hydropathy and aliphatic index (ρ=0.28).

Based on these correlation patterns, three physicochemical properties—charge, hydropathy, and aliphatic index—were selected for subsequent analyses. These descriptors capture complementary electrostatic, hydrophobic, and structural stability-related characteristics that may influence spike protein conformation, intermolecular interactions, and functional evolution, while minimizing redundancy among correlated physicochemical variables. Accordingly, each spike protein sequence was represented as a vector in a three-dimensional physicochemical space (ℝ^3^).

### Spike evolution remains confined to a restricted physicochemical space despite continued sequence diversification

To quantify the divergence of sequence-derived physicochemical vectors from a reference baseline, we calculated the squared Mahalanobis distance. Similar multivariate distance measures have previously been used to characterize viral evolution and sequence divergence (60). We examined the resulting trajectories as functions of both elapsed time and the number of amino acid substitutions, which reached a maximum of 90 in the dataset. The squared Mahalanobis distance increased rapidly during the first two years of the pandemic, subsequently reached a maximum, and then declined (Fig. 1A). A similar trajectory was observed when physicochemical displacement was examined as a function of the number of amino acid substitutions rather than calendar time (Fig. 1B).

**Fig 1.**
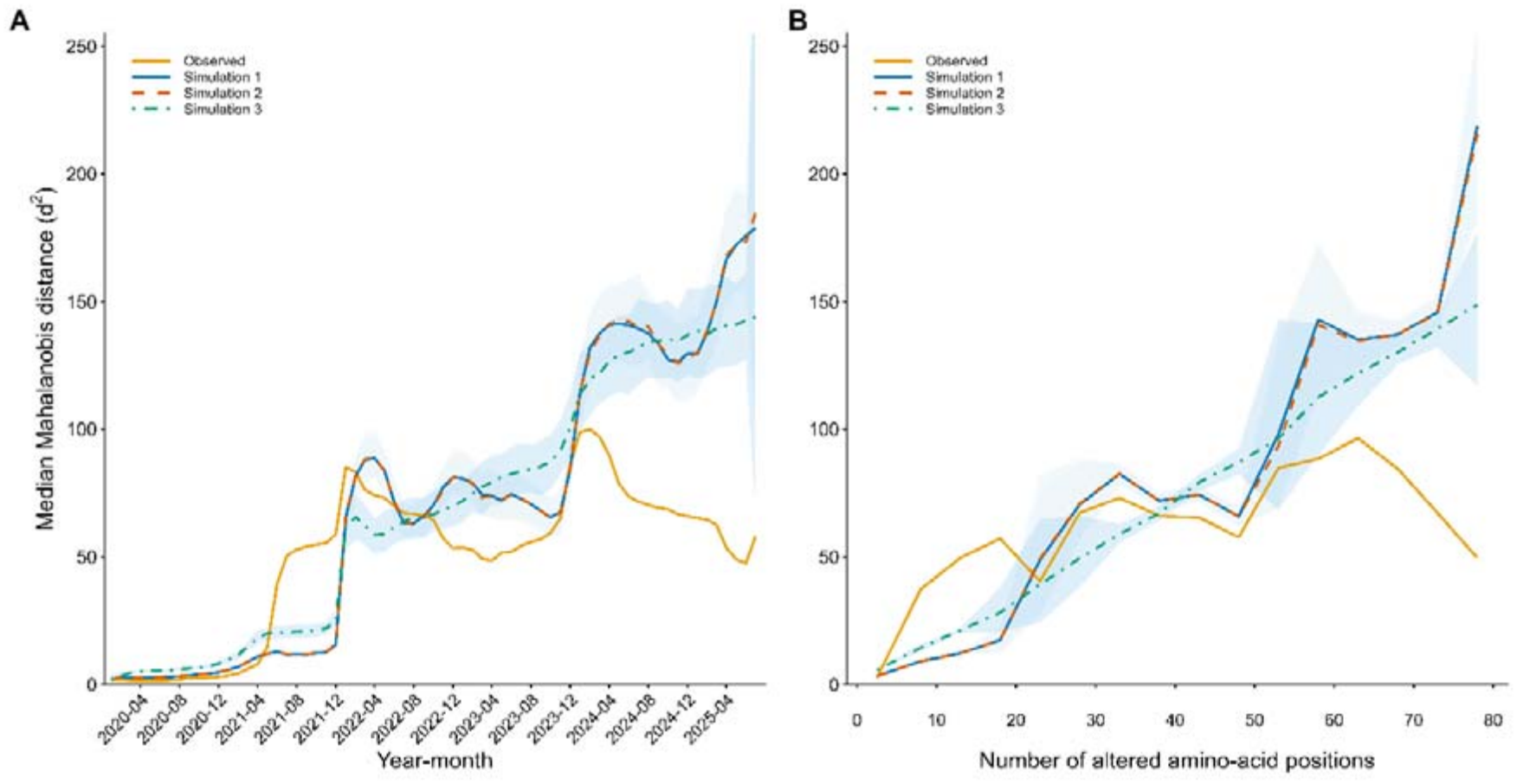
Median squared Mahalanobis distances of observed and simulated physicochemical vectors. (A, B) Displacement of observed sequence-derived and simulation-derived physicochemical vectors from the baseline, plotted against calendar time (A) and the number of variable positions (B). The solid orange line represents the observed median squared Mahalanobis distance (*d*^2^). The solid blue and dashed red lines represent the median *d*^2^ values from the memoryless and memory-based simulations, respectively, in which the empirical sequence-specific constellations of variable positions were retained while amino acid identities were randomized. The dot-dashed green line represents the median *d*^2^ from the whole-spike memoryless simulation, in which the number of randomized positions per simulated sequence was constrained to match the number of distinct variable positions in the corresponding empirical sequence, but their locations were randomized across the entire spike protein. In all three simulation models, amino acids were sampled with equal probability from the 20 standard amino acids. Shaded areas indicate the 95% of Monte Carlo null envelopes, defined by the 2.5^th^ and 97.5^th^ percentiles of the median *d*^2^ distributions across 200 independent realizations of each simulation model at each calendar time point (A) or variable position class (B). The baseline was defined using physicochemical vectors derived from sequences collected during the first three months of the pandemic.

To assess whether these observed patterns could be reproduced under the conditional randomization processes represented by the three null models, we compared the empirical trajectories with those obtained from Monte Carlo simulations. In contrast to the empirical data, all three simulation schemes exhibited a progressive increase in squared Mahalanobis distance, both over time (Fig. 1A) and with increasing numbers of amino acid substitutions (Fig. 1B). Thus, none of the three randomization schemes reproduced the nonmonotonic trajectory observed in the empirical data. These findings suggest that the physicochemical evolution of the spike protein was constrained relative to the evolutionary patterns expected under the specified null models.

Collectively, these results indicate that, despite the continued accumulation of amino acid substitutions, SARS-CoV-2 spike protein evolution remained confined to a restricted region of physicochemical space.

### Clustering identified six phenotypic groups

To investigate the organization of the constrained physicochemical space, protein sequences from the 12 study regions were clustered according to their charge at pH 7.0, hydropathy, and aliphatic index. Parameter optimization based on CTL and ΔCTL, together with assessments of clustering stability, supported the selection of α = 0.02 and *k* = 6 (Supplementary Fig. 1), with an eigenvector ratio restriction factor of 50. At α = 0.02, the six-cluster solution achieved the highest median adjusted Rand index (ARI = 0.747; IQR = 0.060), compared with 0.739 and 0.693 for the five- and seven-cluster solutions, respectively, further supporting the selection of k = 6. Comparison of the candidate solutions also showed that *k* = 6 provided greater overall physicochemical differentiation than *k* = 5, with a median absolute standardized difference of 2.389 versus 1.667, respectively. Although the seven-cluster solution yielded a similar value (2.344), it provided limited additional differentiation. Temporal analysis further indicated that the six-cluster solution offered greater resolution than the five-cluster solution, revealing more pronounced periods of reduced cluster dominance. In contrast, the seven-cluster solution produced broadly similar temporal patterns, with only marginal gains in resolution (Supplementary Fig. 1, C). Collectively, these findings supported the retention of six clusters as an informative representation of the physicochemical organization and temporal dynamics of the spike protein. Cluster 0 represented trimmed observations (outliers) rather than an additional phenotypic cluster and was therefore excluded from subsequent analyses.

Incorporating within-country duplicate sequence frequencies as weights resulted in only minor changes in cluster prevalence (maximum absolute difference, 1.66%), centroid locations (mean Euclidean shift, 0.0265; maximum, 0.0734), and the variance explained by PCA (maximum difference, 0.012). These findings indicate that the inferred physicochemical structure was robust to sequence frequency weighting and was not substantially affected by the removal of duplicate sequences.

Ridge multinomial regression yielded near-zero coefficients (|*β*| ≤ 0.002) for all country and sample-size predictors, indicating limited country-level effects on cluster assignment, despite statistical significance attributable to the large sample size. Consistent with this finding, Cramer’s V indicated only a weak association between geographic origin and cluster membership (0.1873 ± 0.00159), suggesting that geographic origin accounted for only a limited proportion of the observed variation in cluster membership.

Multivariate analysis of variance (MANOVA) demonstrated pronounced physicochemical differentiation among the six clusters (Wilks’ λ = 0.0457, approximate *F*_l5,735206.S_ = 100907.78, p < 2.2 × 10^−16^). Subsequent pairwise comparisons using Pillai’s trace identified significant multivariate differences between all cluster pairs following Benjamini-Hochberg correction (Supplementary Table 5). Cluster stability and separability were further supported by low centroid variability in bootstrap resampling analyses (Supplementary Table 6) and by linear discriminant analysis (LDA), which achieved approximately 96.3% cross-validation accuracy (Supplementary Table 7).

The clustering analysis identified one dominant phenotypic group (cluster 1) comprising 55.3% of all sequences, and four additional major clusters (clusters 2 to 5), which collectively accounted for 41.3% of the unique sequences (Table 2). Cluster 6 represented 1.6 % of the sequences, whereas approximately 2% of unique sequences were assigned to cluster 0, corresponding to outliers excluded from subsequent analyses. Each phenotypic cluster contained sequences representing multiple SARS-CoV-2 variants (Table 2), indicating that genetically distinct viral lineages can converge on similar physicochemical states. Conversely, although more than 80% of the sequences belonging to most variants were assigned to a single phenotypic cluster, smaller proportions were consistently distributed across additional clusters, indicating measurable physicochemical heterogeneity within individual variants (Supplementary Fig. 2).

**Table 2.**
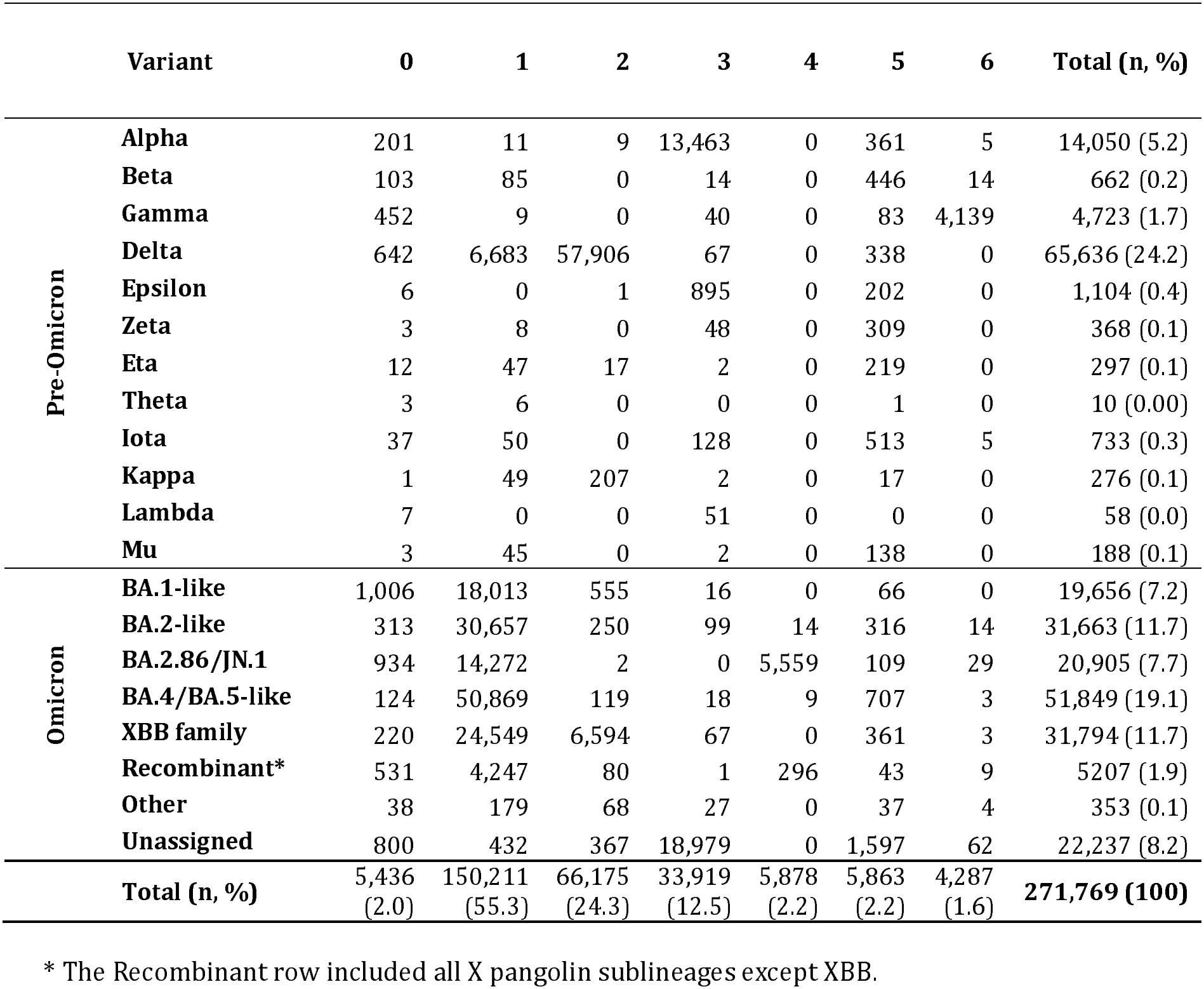
Distribution of SARS-CoV-2 variants across phenotypic clusters defined by three physicochemical properties: charge at pH 7.0, hydropathy, and aliphatic index. Each cell indicates the number of unique spike amino acid sequences belonging to a given genetic variant (rows) and phenotypic cluster (columns).

The six clusters exhibited distinct physicochemical profiles (Supplementary Fig. 3), characterized by differences in parameter distributions and degrees of within-cluster variability (Supplementary Figs. 4 and 5; Supplementary Table 8). Interactive visualizations of these distributions are available at: https://inminz.github.io/sarscov2-spike-cluster-variant-physicochemical-means/ and https://inminz.github.io/sarscov2-spike-variant-physicochemical-means-by-cluster/, respectively.

### PCA revealed an organized physicochemical space

Principal component analysis (PCA) of the standardized physicochemical variables (interactive visualization available at https://inminz.github.io/sarscov2-spike-physicochemical-pca) yielded three principal components (PCs), collectively accounting for 100% of the total variance (Supplementary Fig. 6). PC1 reflected relatively balanced contributions from all three variables, with the aliphatic index contributing slightly more than hydropathy and charge (Table 3). PC2 was primarily driven by charge, followed by hydropathy, whereas the contribution of the aliphatic index was minimal. In contrast, PC3 was characterized by nearly equal contributions from hydropathy and the aliphatic index, with negligible contribution from charge.

**Table 3.** Principal component analysis (PCA) eigenvalues (first row), coefficients, and relative contribution of each physicochemical property to the identified principal components.

| **Variable** | **PC1 coefficient** | **PC1 (%)** | **PC2 coefficient** | **PC2 (%)** | **PC3 coefficient** | **PC3 (%)** |
|---|---|---|---|---|---|---|
| Eigenvalue | 1.313 |  | 0.829 |  | 0.684 |  |
| Charge at pH 7.0 | -0.528 | 27.86 | 0.836 | 69.9 | -0.15 | 2.24 |
| Hydropathy | 0.579 | 33.55 | 0.483 | 23.35 | 0.657 | 43.11 |
| Aliphatic index | 0.621 | 38.59 | 0.26 | 6.76 | -0.739 | 54.65 |

Coefficients are signed eigenvector components; percentages are squared-coefficient contributions.

Projection of the sequences onto the principal component axes revealed distinct separation among most clusters, indicating an organized low-dimensional physicochemical landscape. Rather than being distributed uniformly across the physicochemical space, spike proteins occupied distinct regions corresponding to discrete physicochemical states. This pattern is consistent with the constrained and finite combinatorial nature of amino acid sequence space (GitHub: https://inminz.github.io/sarscov2-spike-physicochemical-pca).

The distribution of clusters across the PC1-PC2 plane primarily reflected differences in charge, with a progression from the negative values characteristic of early (unassigned) and Alpha lineages toward the positive values observed among Omicron variants. In the PC1-PC3 plane, separation was driven predominantly by hydropathy, with clusters 2 and 6 occupying opposite extremes along this axis. Finally, the PC2-PC3 plane highlighted differences associated with charge and aliphatic index. Clusters 1 and 2 exhibited the highest aliphatic index values, whereas clusters 3 and 6 exhibited the lowest.

### Genotypic-phenotypic spatiotemporal dynamics

Distinct temporal relationships between variant entropy and physicochemical cluster entropy were observed across geographic regions (Fig. 2 and Supplementary Figs. 7 and 8). Although the two measures peaked simultaneously during certain periods, extended intervals of decoupling were also evident. Across all regions, normalized genotypic entropy was generally higher than normalized phenotypic entropy, with negative mean ΔH values observed in 10 of the 12 regions (Supplementary Table 9). This pattern suggests that extensive diversification of amino acid sequence constellations was accompanied by considerably more limited diversification of physicochemical phenotypes. Nevertheless, the difference between phenotypic and genotypic entropy (ΔH) fluctuated over time across geographic regions (Fig. 3).

**Fig 2.**
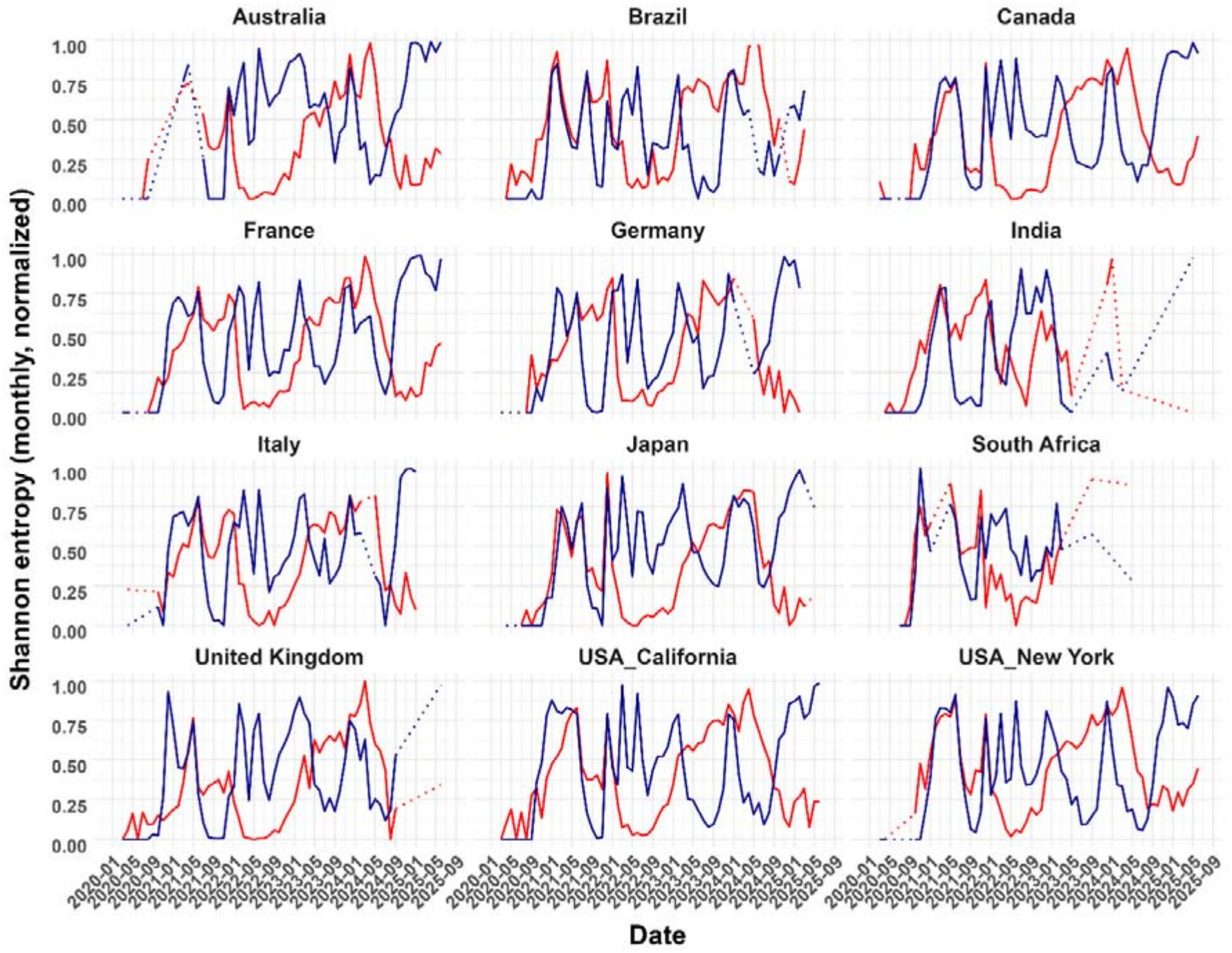
Temporal dynamics of normalized Shannon entropy of genotypic and phenotypic diversity across 12 geographic regions. Genotypic diversity (dark blue) was defined using the WHO–Pango key classification, whereas phenotypic diversity (red) was defined based on physicochemical clusters. Entropy was estimated monthly using equal-sample-size (N) subsampling. Dotted line segments indicate periods with insufficient sample sizes for reliable entropy estimation.

**Fig 3.**
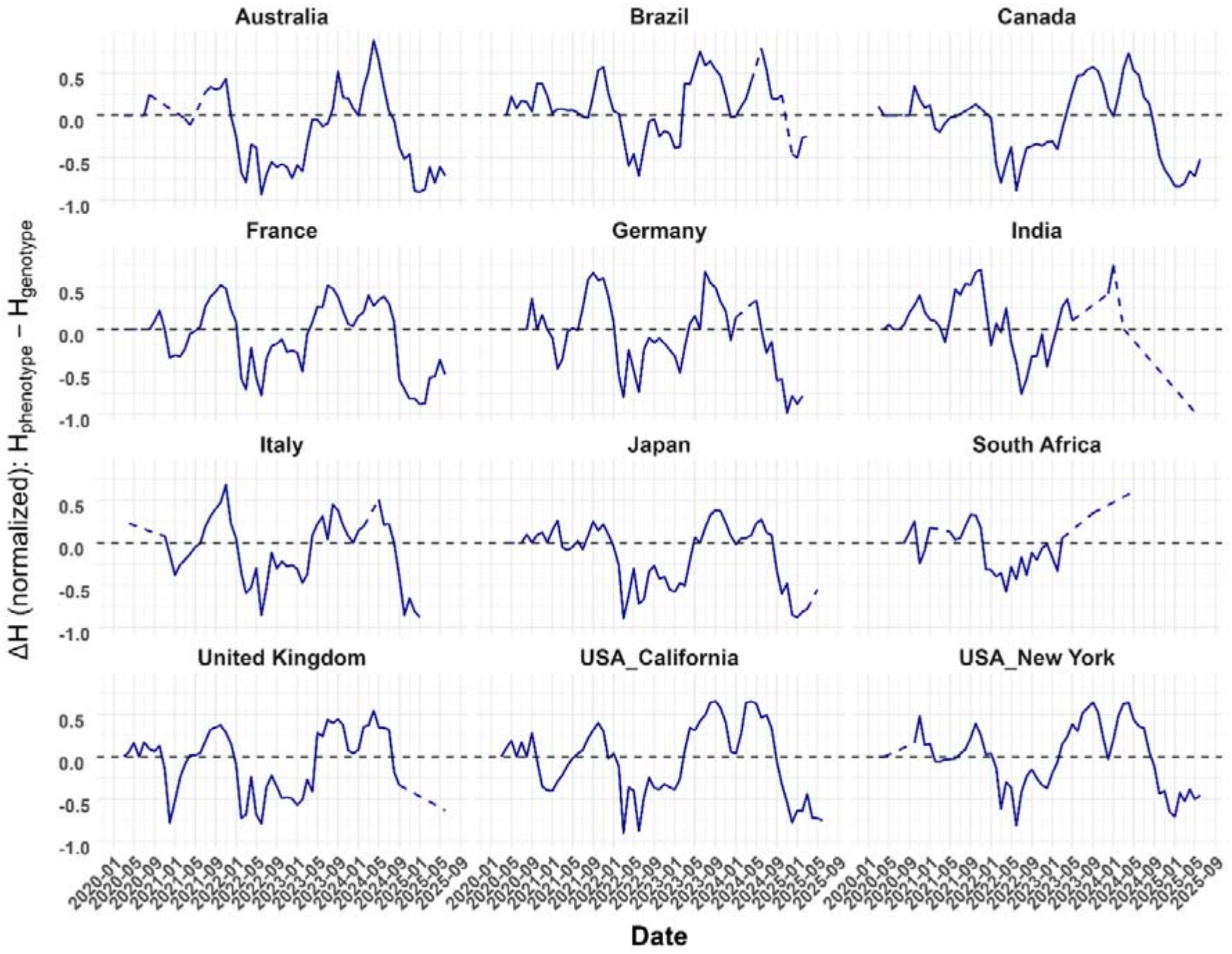
Temporal dynamics of the difference between normalized phenotypic and genotypic Shannon entropy (ΔH = phenotypic – genotypic) across 12 geographic regions. Positive values indicate higher phenotypic than genotypic entropy, whereas negative values indicate higher genotypic than phenotypic entropy. Entropy was estimated monthly using equal-sample-size (N) subsampling. Dashed line segments indicate periods with insufficient sample sizes for reliable entropy estimation.

Measures of decoupling, including the mean, median, maximum, and 95^th^ percentile of |ΔH|, revealed substantial geographic heterogeneity (Supplementary Table 9). Mean absolute decoupling ranged from 0.228 to 0.411, with maximum monthly values reaching 0.983. The proportion of months during which genotypic entropy exceeded phenotypic entropy ranged from 31.0% to 64.2%, further indicating regional differences in the temporal relationship between genotypic and phenotypic diversity.

Generalized additive model (GAM) analysis demonstrated a highly significant nonlinear temporal pattern in the magnitude of phenotype-genotype decoupling (|*ΔH*|; effective degrees of freedom [edf] = 7.985, F = 23.02; p = 2 × 10^-16^). LOESS regression revealed broadly similar temporal patterns across geographic regions, despite regional differences in the duration and magnitude of decoupling throughout the pandemic (Supplementary Fig. 9). However, pairwise Wilcoxon tests detected no statistically significant differences in the magnitude of decoupling among geographic regions after correction for multiple comparisons (minimum adjusted p = 0.33).

## Discussion

The principal finding of this study is that extensive SARS-CoV-2 spike sequence diversification remained confined to a limited physicochemical space. Although amino acid substitutions substantially altered spike sequence composition, they did not produce the progressively increasing physicochemical displacement expected under empirically conditioned randomization models designed to preserve key features of the observed evolutionary process. Instead, the observed squared Mahalanobis distances indicated that evolutionary trajectories remained within relatively narrow bounds, consistent with physicochemical constraints on spike diversification.

Notably, the physicochemical trajectories generated by the memoryless and memory-based models, both of which were conditioned on the observed number and location of variable positions, were highly similar. Thus, introducing temporal dependence by retaining randomized amino acid outcomes at recurring variable positions had little effect on physicochemical displacement relative to independent randomization. These findings suggest that temporal memory alone is insufficient to explain the constrained physicochemical trajectories of naturally evolving spike sequences. In contrast, the memoryless model, in which amino acids were randomized across the entire spike sequence while preserving only the observed number of variable positions, produced substantially greater physicochemical displacement. This contrast highlights the importance of the positional constraints on sequence variation and the contribution of the largely conserved protein background to the maintenance of spike physicochemical properties.

The greater physicochemical displacement observed under unrestricted positional randomization further suggests that naturally variable positions may represent evolutionarily permissive sites at which amino acid variation can explore a broader range of physicochemical properties without necessarily compromising structural viability. This interpretation is consistent with recent evidence that structural and thermodynamic constraints restrict the sequence space accessible to the SARS-CoV-2 spike protein (61).

Together, these findings indicate that spike evolution is constrained at the level of amino acid constellations, restricting sequence diversification to a bounded physicochemical space. Consistent with this interpretation, clustering and PCA revealed that spike variation occupied a low-dimensional region of physicochemical space partitioned into six distinct clusters. These clusters represent recurrent physicochemical phenotypes arising from a restricted subset of amino acid combinations. The organization of spike diversity is consistent with previous work (15), which demonstrated that physicochemical representations of spike sequences can capture structural characteristics associated with SARS-CoV-2 lineages.

Within this framework, the phenotypic clusters represent recurrent physicochemical states rather than regions uniquely associated with individual variants. For example, genetically divergent variants, including early variants such as Alpha and Delta and later Omicron sublineages, occupied similar regions of physicochemical space despite substantial differences in the number and composition of amino acid substitutions and in their times of emergence. These observations suggest that distinct evolutionary trajectories can converge on similar physicochemical configurations despite extensive genetic divergence.

Moreover, individual variants exhibited measurable physicochemical heterogeneity, characterized by a dominant phenotype accompanied by less frequent alternative configurations occupying distinct regions of physicochemical space. This observation is consistent with previous finding of substantial genotypic heterogeneity within individual SARS-CoV-2 variants, in which numerous amino acid constellations were identified beyond their canonical variant-associated signatures (62). Collectively, these findings demonstrate that distinct amino acid constellations within a single variant can correspond to distinguishable physicochemical phenotypes. Consequently, cluster assignment appears to depend primarily on the overall physicochemical configuration of the spike protein rather than on variant designation, genetic distance, or chronological order of emergence.

Overall, these observations support a many-to-one genotype-phenotype mapping (63–65), whereby diverse amino acid configurations converge on a limited set of recurrent physicochemical phenotypes. In this context, the experimentally observed mutational tolerance of the SARS-CoV-2 spike protein (66, 67), including its capacity to accumulate amino acid substitutions while maintaining or enhancing ACE2 binding and facilitating immune escape (66), may be interpreted as diversification within a bounded physicochemical space. This interpretation provides a framework for understanding how substantial sequence variation can occur without necessarily producing proportionate changes in the physicochemical properties captured by our descriptors.

We next examined the spatiotemporal dynamics of physicochemical space occupancy using Shannon entropy. Our analysis revealed dynamic genotype-phenotype decoupling throughout the pandemic. Periods characterized by ΔH > 0 reflected transient increases in phenotypic diversity relative to genotypic diversity, consistent with a broader distribution of circulating amino acid constellations across physicochemical states. Conversely, prolonged periods characterized by ΔH < 0 indicated that extensive genetic diversification was accompanied by comparatively limited physicochemical diversification, suggesting that multiple genetically distinct amino acid constellations converged on a restricted set of physicochemical phenotypes.

This pattern is consistent with the hypothesis that only a subset of theoretically possible amino acid constellations is evolutionarily compatible and that these constellations represent alternative evolutionary pathways (62). Our findings further suggest that this restricted compatibility may be reflected not only in sequence composition but also in physicochemical properties, with circulating amino acid constellations occupying a relatively restricted region of physicochemical space. Conversely, constellations that fail to persist during viral evolution may be less likely to attain or maintain physicochemical configurations compatible with continued viral propagation. However, because the present analysis is observational, the relationship between physicochemical configurations and evolutionary persistence remains an inference rather than a direct demonstration of functional compatibility.

Our GAM analysis revealed a significant nonlinear temporal pattern in the magnitude of genotype-phenotype decoupling across the populations examined. Despite differences in the duration and magnitude of these patterns, broadly similar temporal trajectories were observed across countries. This similarity suggests that, although the distribution of amino acid constellations varied among populations, spike evolution repeatedly occupied similar physicochemical states. Such convergence is consistent with shared selective constraints associated with spike function and adaptation to human hosts, although the present analysis cannot establish the relative contributions of specific selective pressures. In this context, transitions between periods of genotype–phenotype coupling and decoupling may reflect alternating phases of sequence-space exploration and selective filtering. Viral evolution can thus be conceptualized as the exploration of sequence space, with natural selection potentially favoring a subset of amino acid constellations that maintain physicochemical properties compatible with infection and transmission. Population-specific characteristics may, in turn, influence how this physicochemical space is explored.

Several limitations should be considered when interpreting these findings. First, this study was based on observational data, and the observed patterns of variation may have been influenced by multiple uncontrolled factors. Although the large sample size and application of statistical approaches designed to characterize evolutionary patterns strengthen the analysis, residual sources of variability and potential sampling biases cannot be excluded. Second, the physicochemical space examined here was defined by three key properties of the spike protein. Although these descriptors captured major axes of variation and discriminated among the observed physicochemical phenotypes, additional structural, biophysical, or functional descriptors relevant to virus-host interactions may reveal further dimensions of evolutionary constraints and refine the organization of this space. Third, the structural and functional consequences of variation along these physicochemical dimensions were not directly evaluated. Accordingly, the inferred physicochemical constraints should not be equated with experimentally established structural or functional restrictions. Nevertheless, the central finding remains that SARS-CoV-2 spike sequence variation was confined to a limited physicochemical space despite extensive diversification over the course of the pandemic.

The physicochemical space characterized here should not, nonetheless, be regarded as an immutable property of SARS-CoV-2. Rather, it represents an empirically reconstructed evolutionary framework that may reflect the cumulative effects of the selective pressures arising from host-virus interactions. This interpretation raises the possibility of using recurrent viral physicochemical states to inform reverse evolutionary inference. Instead of directly characterizing the physicochemical properties of host cellular environments, future studies could investigate whether recurrent viral physicochemical configurations provide indirect evolutionary signatures of the constraints associated with successful infection. Such an approach could generate hypotheses about the physicochemical landscape shaping viral adaptation, although independent structural, biochemical, and functional validation would be required to establish the underlying mechanisms.

Whether viruses with similar cell-entry mechanisms occupy comparable physicochemical spaces, and whether analogous landscapes emerge during the evolution of other rapidly evolving viruses, remain important questions for future research. The quantitative framework presented here, incorporating measures of physicochemical space occupancy, phenotypic diversity, and genotype–phenotype decoupling, may facilitate comparative analyses across diverse animal virus systems. More broadly, the evolutionary principles underlying physicochemical constraints may also be relevant to other host-virus systems, including plant viruses. Thus, the physicochemical space identified in this study may be viewed not as a fixed characteristic of SARS-CoV-2 but as an emergent evolutionary framework linking sequence diversity to physicochemical variation under changing selective environments.

## Acknowledgements

Partial results of this work were presented at the ECV2025 conference, Dubrovnik, Croatia (Abstract No. 423). We gratefully acknowledge all data contributors, including the authors and originating laboratories responsible for obtaining the specimens, as well as the submitting laboratories responsible for generating genetic sequence data and associated metadata and sharing these resources through the GISAID Initiative, upon which this research is based.

## Funding

JAH was supported by the Medical Research Council (MRC) MR/W005611/1 to G2P-UK: a national virology consortium to address phenotypic consequences of SARS-CoV-2 genomic variation and the MRC MR/Y004205/1 to the G2P2 virology consortium: keeping pace with SARS-CoV-2 variants, providing evidence to vaccine policy, and building agility for the next pandemic.

## Notes

### Competing Interest Statement

The authors have declared no competing interest.

