## Supplementary Figures for "Extensive SARS-CoV-2 spike sequence diversification converges on a bounded, low-dimensional physicochemical space"

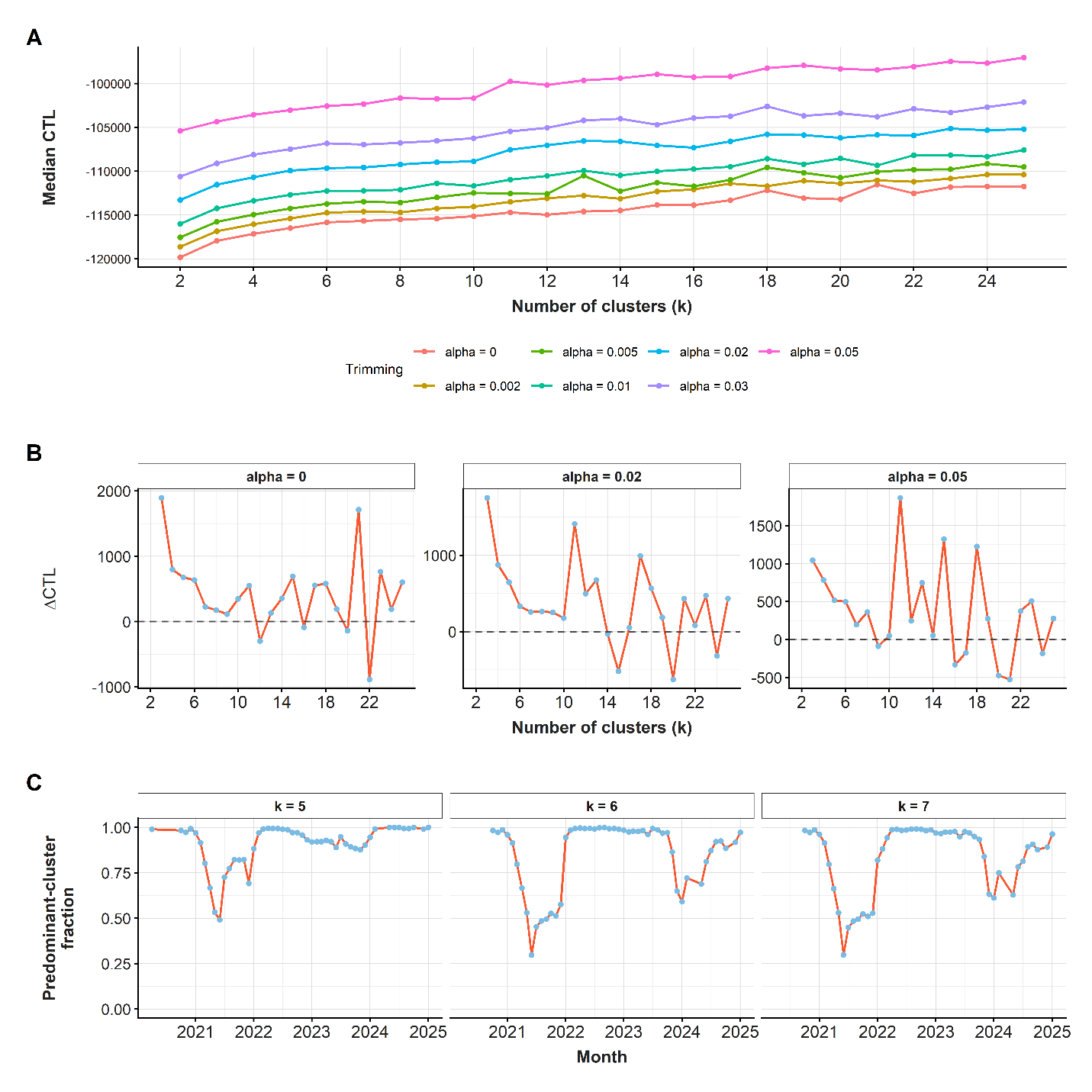

**Supplementary Fig. 1. Selection of the trimming proportion (*α*) and number of clusters (*k*) using the classification trimmed likelihood (CTL) criterion.** (A) CTL as a function of the number of clusters (*k*) across the seven trimming proportions (α = 0, 0.002, 0.005, 0.01, 0.02, 0.03, and 0.05). (B) Successive marginal changes in CTL (ΔCTL) with increasing k across the seven trimming proportions. The horizontal dashed line indicates ΔCTL = 0, distinguishing positive from negative increments in CTL. (C) Temporal evolution of the predominant-cluster fraction for solutions with k = 5, 6, and 7 clusters. The six-cluster solution revealed more pronounced reductions in cluster dominance than the five-cluster solution, indicating improved temporal resolution, particularly during late 2023-2025.

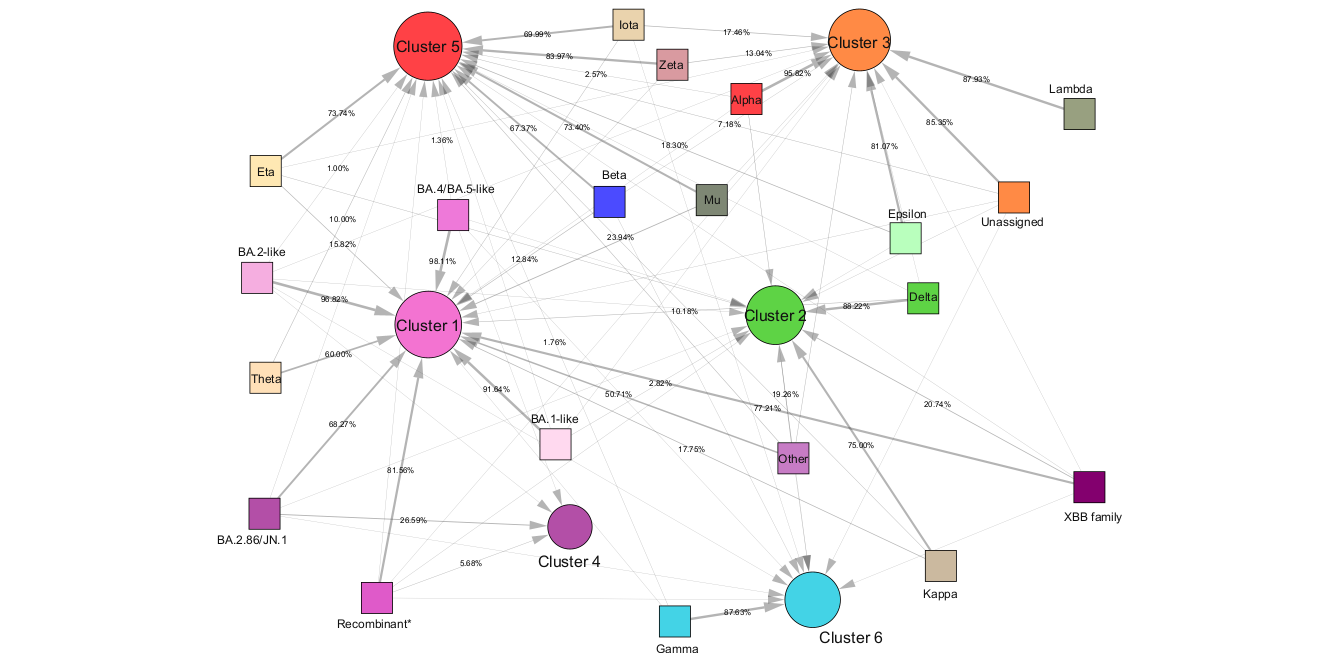

**Supplementary Fig. 2**. **Network representations of the associations between WHO/Pango variants and phenotypic clusters.** Edge thickness is proportional to the percentage of sequences assigned to each phenotypic cluster, reflecting the relative strength of the association between variants and phenotypic clusters. “Recombinant” refers to recombinant lineages other than those belonging to the XBB family, as described in Supplementary Table 2.

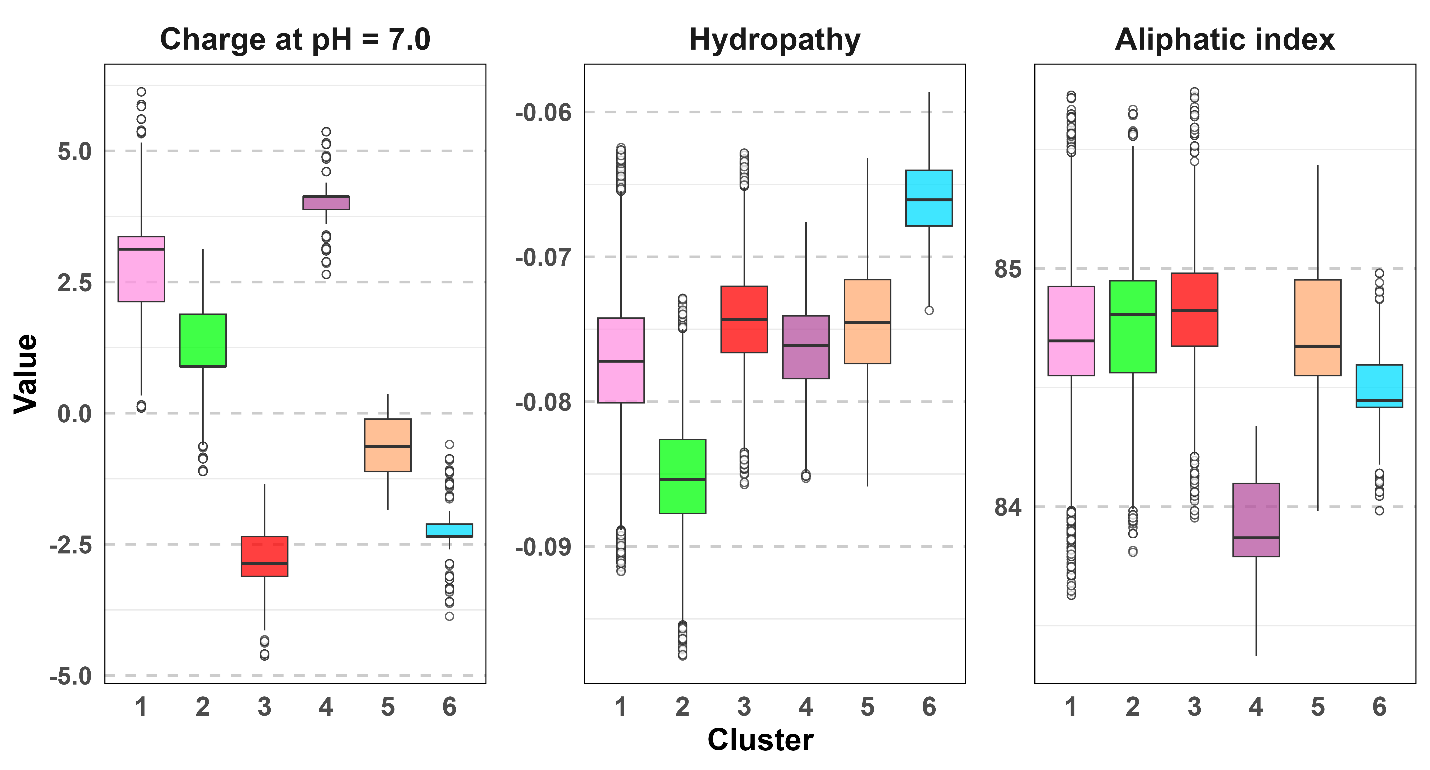

**Supplementary Fig. 3.** **Distributions of physicochemical properties across phenotypic clusters.** Box plots illustrate the distributions of charge at pH 7.0, hydropathy, and aliphatic index across the identified phenotypic clusters. The clusters display distinct physicochemical profiles, although some overlap is observed for individual properties. Notably, clusters with similar values for two properties could be distinguished by differences in the third, highlighting the complementary contribution of all three descriptors to cluster discrimination.

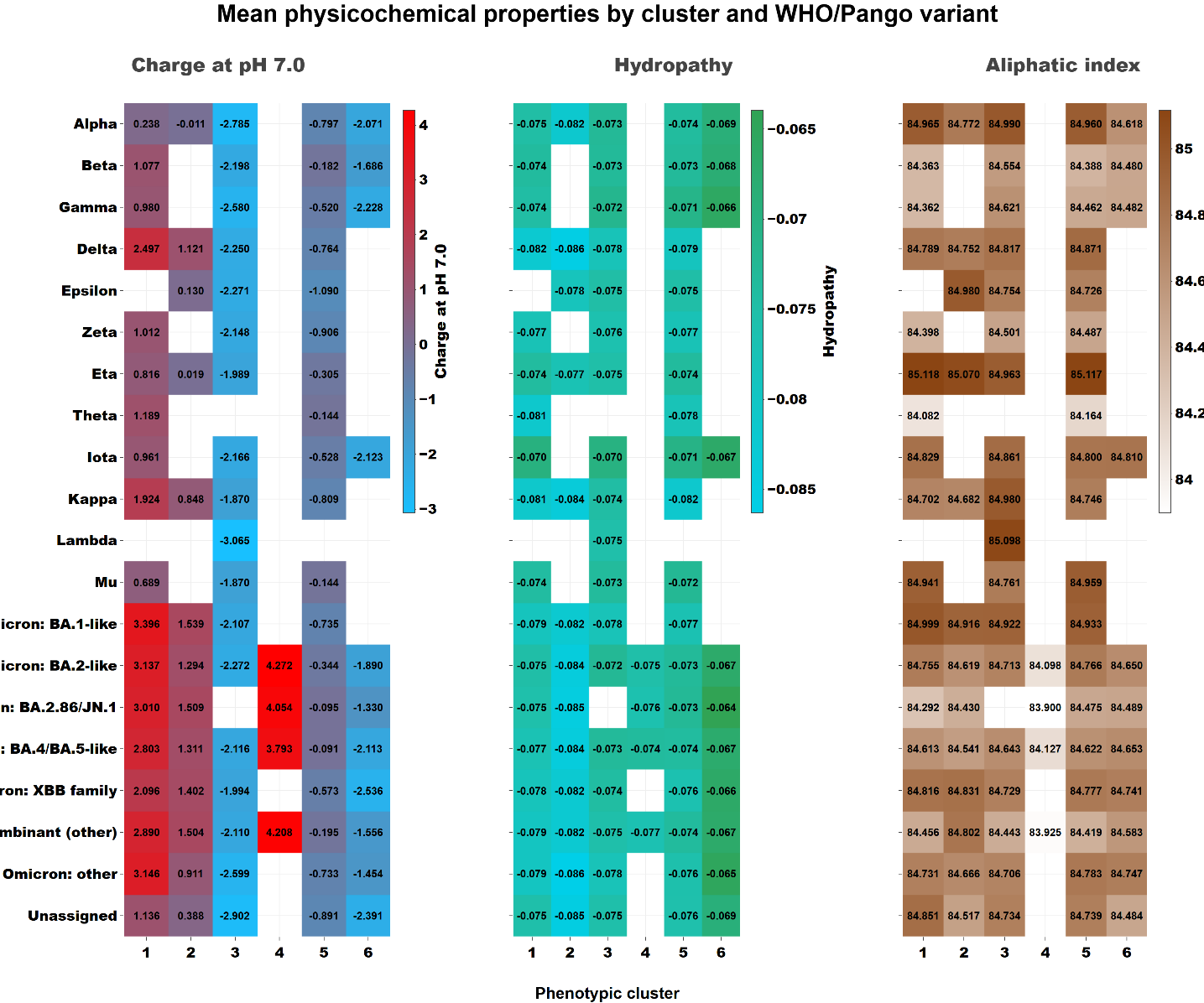

**Supplementary Fig. 4**. **Mean physicochemical properties across WHO/Pango variants and phenotypic clusters.** Mean values for charge at pH 7.0, hydropathy, and aliphatic index are presented for each WHO/Pango variant across the six phenotypic clusters. The sample size (n) for each variant–cluster combination is indicated alongside the corresponding mean values. An interactive version of the figure is available at <https://inminz.github.io/sarscov2-spike-cluster-variant-physicochemical-means/>.

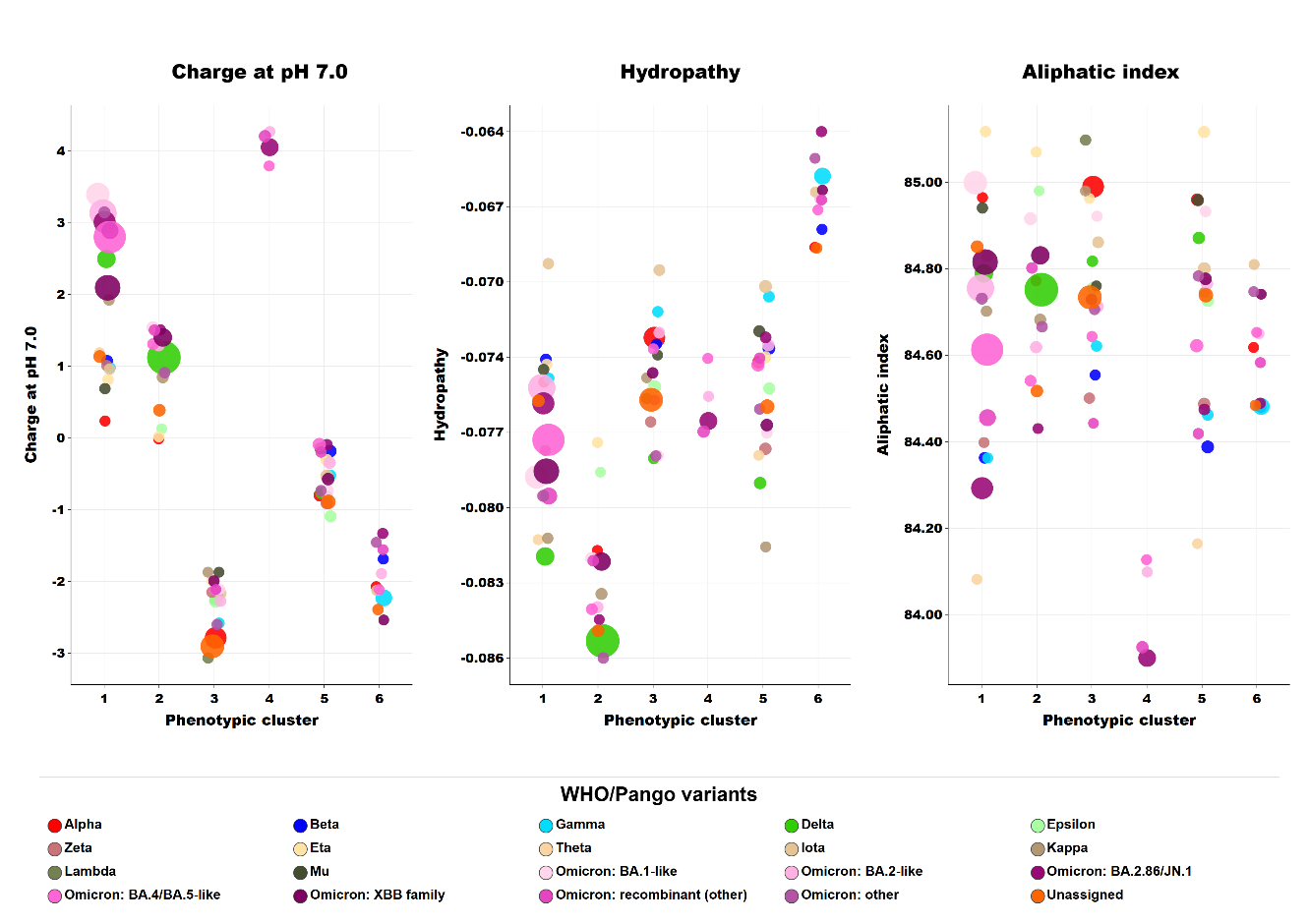

**Supplementary Fig. 5.** **Variant-level mean physicochemical properties within the six phenotypic clusters.** Each point represents the mean physicochemical profile of a WHO/Pango variant within a given phenotypic cluster. Point color denotes the corresponding WHO/Pango variant, while point size is proportional to the number of sequences contributing to the respective mean. An interactive version of the figure is available at <https://inminz.github.io/sarscov2-spike-variant-physicochemical-means-by-cluster/>.

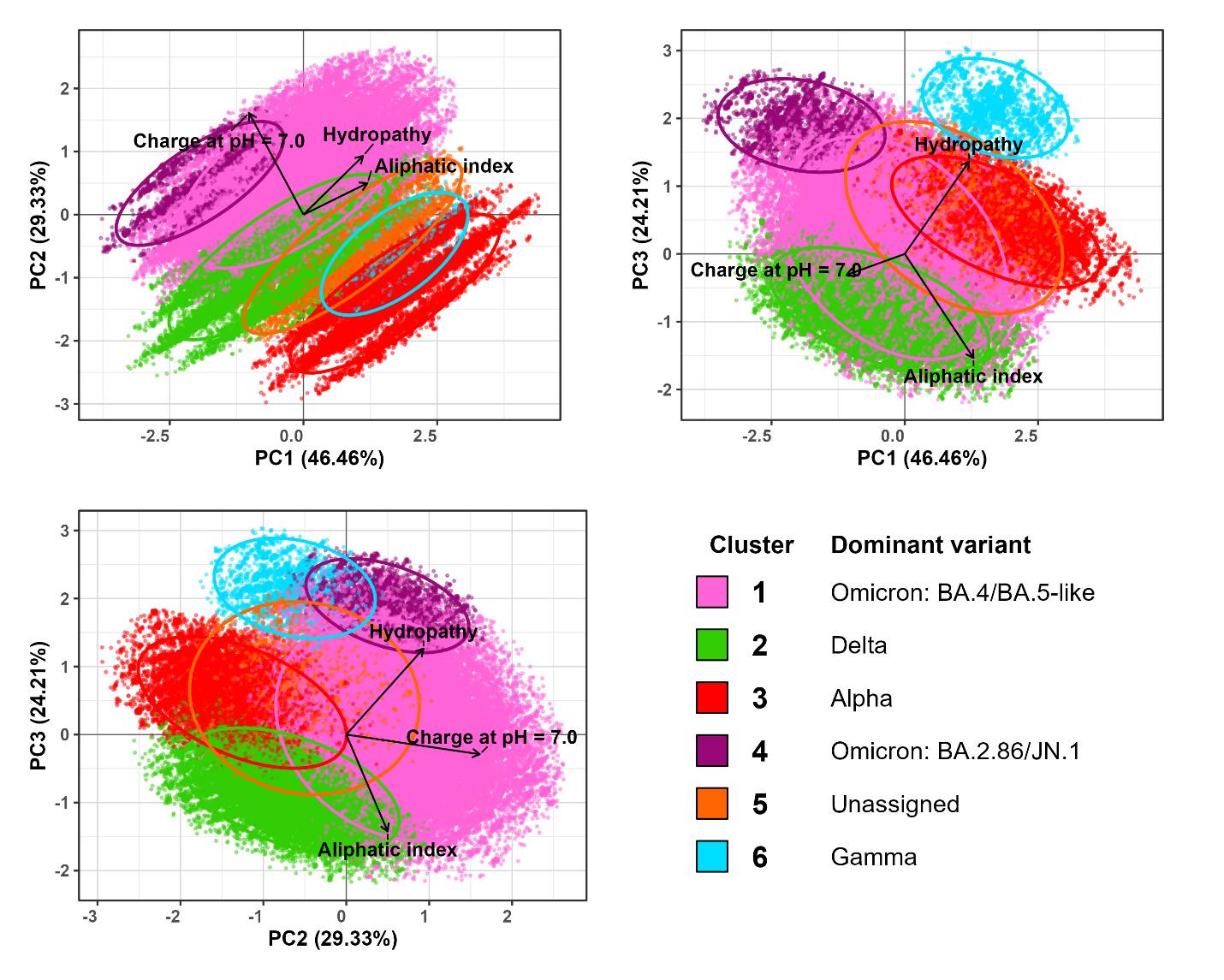

**Supplementary Fig. 6.** **Principal component analysis (PCA) of the six phenotypic clusters based on physicochemical properties.** The three panels show the distribution of sequences in pairwise projections of the principal components (PC1-PC2, PC1-PC3, and PC2-PC3), which together account for 100% of the total variance. Clusters were defined using three physicochemical features: charge at pH 7.0, hydropathy, and aliphatic index. Colors indicate the most prevalent WHO/Pango variant within each cluster. Cluster 3 is colored according to Alpha, the second most frequent variant in this cluster after Unassigned. Vectors represent variable loadings, indicating the direction and relative contribution of each physicochemical property to the principal component space. Ellipses delineate the 95% confidence regions of the cluster distributions, illustrating within-cluster variability. An interactive visualization of the physicochemical space is available at <https://inminz.github.io/sarscov2-spike-physicochemical-pca/>.

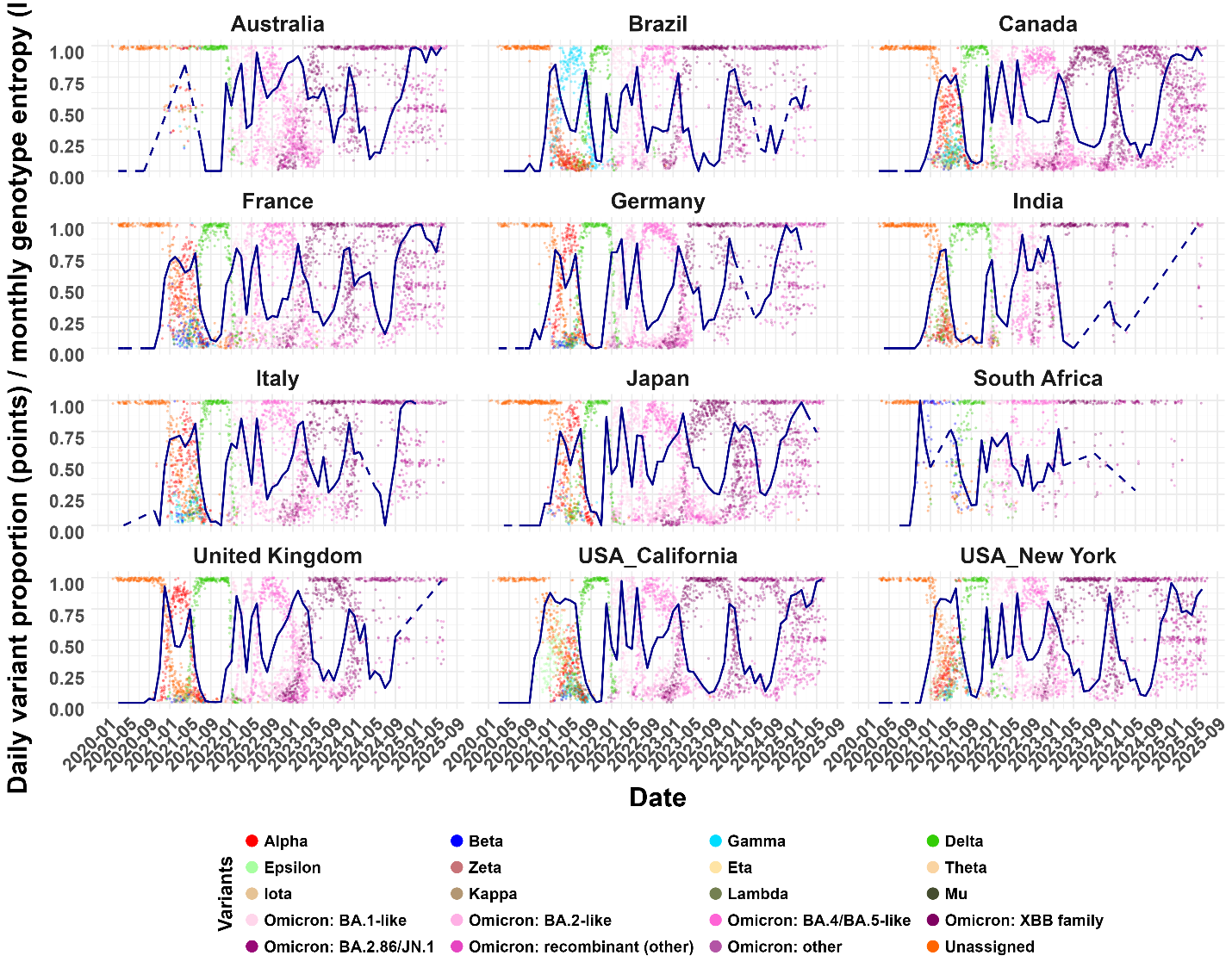

**Supplementary Fig. 7.** **Temporal distribution of WHO-designated SARS-CoV-2 variants across countries during the pandemic.** The figure illustrates the temporal distribution of viral variants, classified according to WHO designations, across different geographic regions. Each dot represents an individual sequence, with vertical jitter applied to reduce overplotting and improve visualization. The blue line represents the LOESS-smoothed monthly Shannon entropy, reflecting temporal changes in variant diversity. Regional differences in variant prevalence over time highlight patterns of variant emergence, dominance, and replacement, capturing changes in viral diversity and epidemiological dynamics throughout the pandemic. Dashed line segments denote periods with insufficient sample sizes for reliable entropy estimation. For visualization purposes, the entropy curves were rescaled to the range of variant proportions.

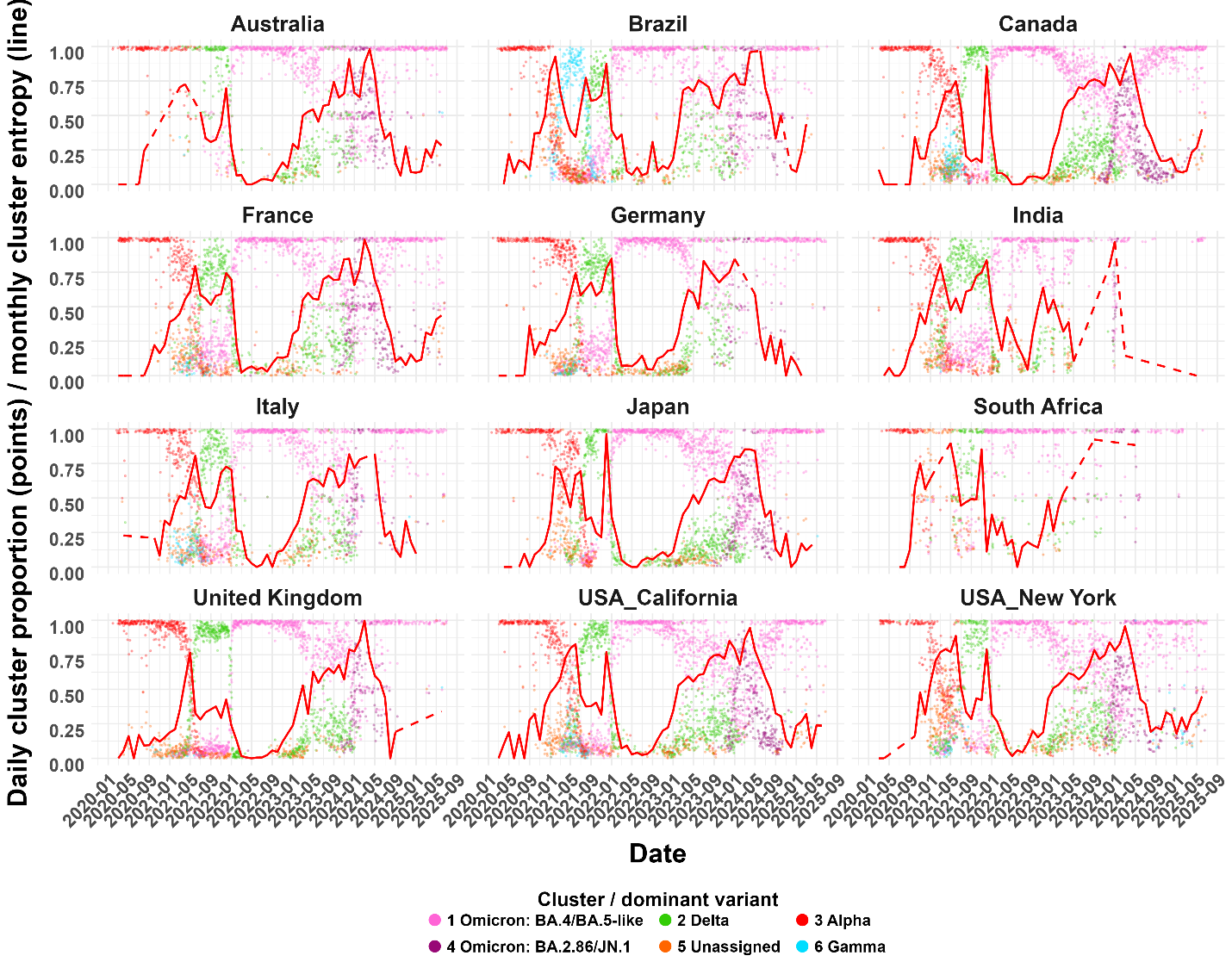

**Supplementary Fig. 8.** **Temporal distribution of physicochemical clusters across geographic locations throughout the pandemic.** Colored dots represent individual sequences assigned to one of the six physicochemical clusters and are plotted according to sampling date. Vertical jitter was applied to reduce overplotting and improve visualization. The red line represents the LOESS-smoothed monthly Shannon entropy, reflecting temporal changes in cluster diversity. Regional differences in the temporal prevalence of physicochemical clusters highlight patterns of cluster emergence, persistence, and replacement throughout the pandemic. Dashed line segments denote periods with insufficient sample sizes for reliable entropy estimation. For visualization purposes, the entropy curves were rescaled to the range of cluster proportions.

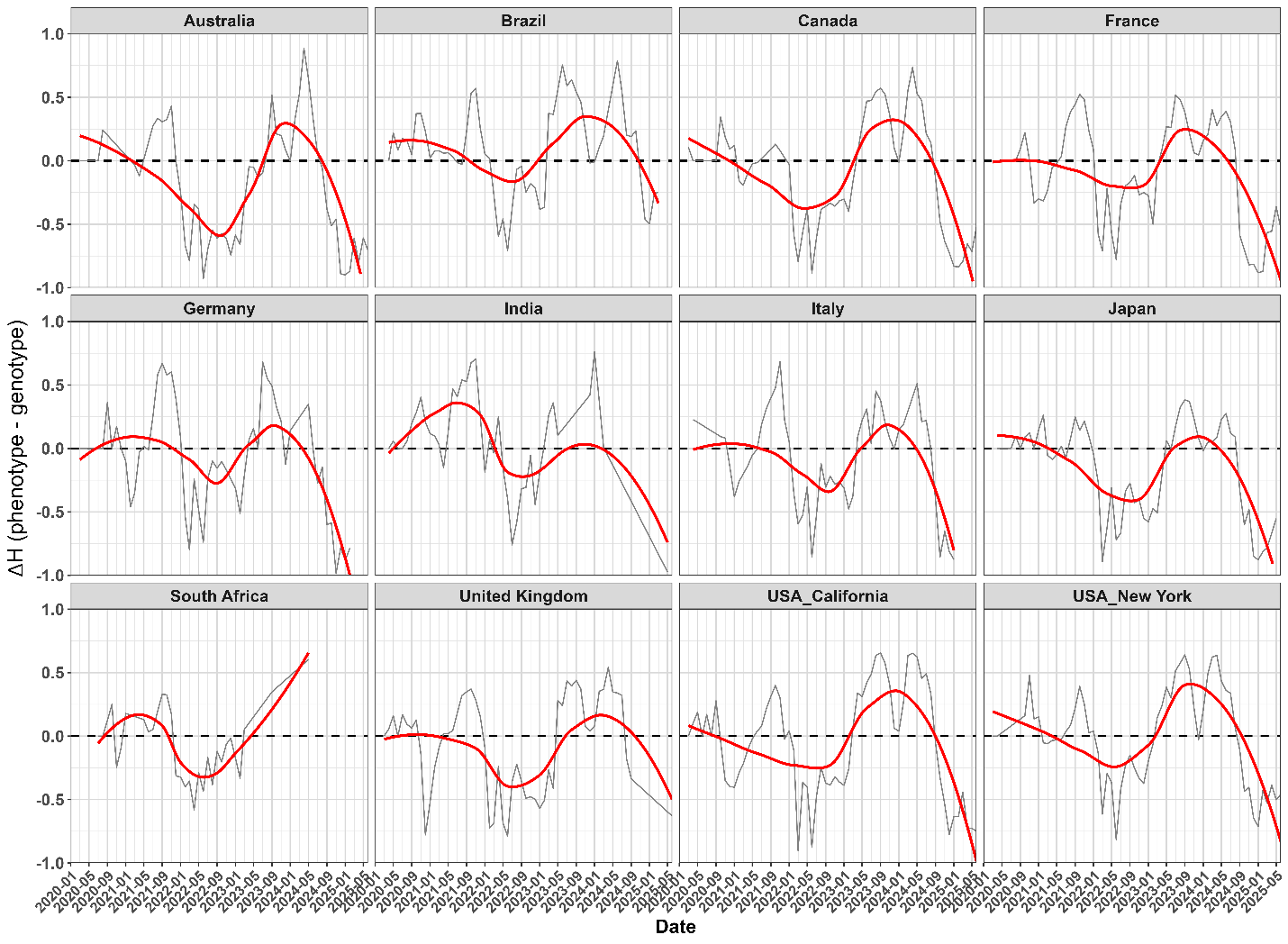

**Supplementary Fig. 9**. **Temporal trajectories of genotype-phenotype decoupling (ΔH = H_phenotype_-H_genotype_) across the 12 geographic regions.** Grey lines represent empirical monthly values, whereas red lines indicate LOESS-smoothed temporal trajectories, highlighting the overall trends within each region. Positive ΔH values indicate that phenotypic diversity exceeds genotypic diversity, while negative values indicate greater genotypic diversity relative to phenotypic diversity.
