## Supplementary Tables for "Extensive SARS-CoV-2 spike sequence diversification converges on a bounded, low-dimensional physicochemical space"

**Supplementary Table 1.** **Workflow for dataset preprocessing and filtering.**

| **Process** | **Description** | **Number of records** | **% of initial dataset** | **% of unique dataset** |
| --- | --- | --- | --- | --- |
| — | All retrieved SARS-CoV-2 spike protein records | 17,113,342 | 100 | — |
| Exclusion | Invalid amino acid sequences | 6,611,562 | 38.63 | — |
| Exclusion | Non-human host sequences | 8,234 | 0.05 | — |
| Exclusion | Sequences shorter than the predefined cutoff (1,263 amino acids) | 192,565 | 1.13 | — |
| Exclusion | Records with ambiguous country annotation | 3,066 | 0.02 | — |
| **Total excluded** |  | 6,815,427 | 39.83 | — |
| **Total after exclusion** |  | 10,297,915 | 60.17 |  |
| Removal | Duplicate sequences (within country) | 9,632,887 | 56.29 | — |
| Result | Unique sequences of all countries | 665,028 | 3.89 | — |
| Result | Sequences retained after geographic-level selection | **272,985** | 1.60 | — |
| Exclusion | Records lacking variant labels | 1,216 | 0.01 | — |
|  | **Dataset used for clustering** | **271,769** | 1.59 | 100.00 |
| Exclusion | Records with invalid dates | 5,338 | 0.03 | 1.96 |
|  | **Dataset used for simulation** | 266,431 | 1.56 | 98.04 |
| Exclusion | Records assigned to Cluster 0 | 5,436 | 0.03 | 2.00 |
|  | **Dataset used for PCA** | 266,333 | 1.56 | 98 |
| Exclusion | Records predating variant emergence | 3,951 | 0.02 | 1.45 |
| Exclusion | Remaining records with invalid dates | 1,146 | 0.01 | 0.42 |
|  | **Dataset used for entropy and decoupling** | 261,236 | 1.53 | 96.12 |

**Supplementary Table 2. Classification of SARS-CoV-2 variants, including WHO-designated labels, dates of first and last detection, geographic origin of the earliest recorded isolate, representative GISAID accession IDs, and corresponding Pango lineage groups.**

| **No.** | **Who name** | **First detection** | **Last detection** | **First place** | **First EPI** | **Last EPI** | **Pango group** |
| --- | --- | --- | --- | --- | --- | --- | --- |
| 1 | Alpha | 01-09-20 | 04-03-24 | Brazil / Minas Gerais | EPI_ISL_19730855 | EPI_ISL_19026222 | B.1.1.7 |
| 2 | Beta | 08-10-20 | 18-04-23 | South Africa / Eastern Cape | EPI_ISL_712081 | EPI_ISL_17796598 | B.1.351; B.1.351.2; B.1.351.3; B.1.351.5 |
| 3 | Gamma | 22-12-20 | 04-01-22 | Dominican Republic | EPI_ISL_9671798 | EPI_ISL_17298323 | P.1; P.1.1; P.1.14; P.1.15; P.1.16; P.1.7.1; P.1.10; P.1.13; P.1.2; P.1.17; P.1.7; P.1.4; P.1.12; P.1.17.1; P.1.11; P.1.10.1; P.1.3; P.1.6; P.1.8; P.1.9; P.1.12.1 |
| 4 | Delta | 03-12-20 | 14-06-24 | Burkina Faso / Centre | EPI_ISL_4255127 | EPI_ISL_19669350 | AY.36; AY.23; B.1.617.2; AY.44; AY.4.2.1; AY.5; AY.100; AY.45; AY.4; AY.25.1; AY.46.6; AY.121.1; AY.43; AY.79; AY.107; AY.20; AY.75; AY.25; AY.26; AY.122; AY.34.1; AY.113; AY.112; AY.124; AY.126; AY.98.1; AY.42; AY.23.2; AY.9.2; AY.85 |
| 5 | Epsilon | 08-10-20 | 29-11-21 | USA / California | EPI_ISL_5695779 | EPI_ISL_11248924 | B.1.429; B.1.427; B.1.429.1 |
| 6 | Zeta | 31-12-20 | 20-05-22 | Canary Islands / Gran Canaria | EPI_ISL_18051052 | EPI_ISL_15436498 | P.2 |
| 7 | Eta | 07-12-20 | 17-12-22 | Cote d’Ivoire / Abidjan | EPI_ISL_9893271 | EPI_ISL_19236220 | B.1.525 |
| 8 | Theta | 24-01-21 | 17-02-22 | Hong Kong | EPI_ISL_1660475 | EPI_ISL_12996407 | P.3 |
| 9 | Iota | 02-12-20 | 20-12-21 | USA / Delaware | EPI_ISL_1137005 | EPI_ISL_12150484 | B.1.526 |
| 10 | Kappa | 05-12-20 | 06-08-21 | India / Maharashtra | EPI_ISL_1360304 | EPI_ISL_4193096 | B.1.617.1 |
| 11 | Lambda | 17-12-20 | 15-11-22 | Peru / Lima | EPI_ISL_8889806 | EPI_ISL_16073474 | C.37; C.37.1 |
| 12 | Mu | 21-01-21 | 07-05-24 | Venezuela / Falcon | EPI_ISL_3507370 | EPI_ISL_19197056 | B.1.621; BB.1; B.1.621.1; BB.2; B.1.621.2 |
| 13 | Omicron: BA.1-like | 04-11-21 | 09-01-25 | Austria / Vienna | EPI_ISL_10741552 | EPI_ISL_19713832 | BA.1; BA.1.17.2; BA.1.17; BA.1.16; BA.1.18; BA.1.1; BA.1.1.18; BA.1.21; BA.1.15.1; BA.1.15; BA.1.13; BA.1.14; BA.3; BA.1.1.14; BA.1.19; BA.1.20; BA.1.12; BA.1.1.11; BA.1.4; BA.1.1.3; BA.1.1.7; BA.1.1.1; BA.1.14.1; BA.1.9; BA.1.1.8; BA.1.8; BA.1.1.2; BA.1.15.2; BD.1; BA.1.1.15 |
| 14 | Omicron: BA.2.86/JN.1 | 24-01-22 | 05-07-25 | Slovakia / Zilinsky kraj | EPI_ISL_19546665 | EPI_ISL_20076243 | KP.1.1.3; JN.1.1; KP.3.1.1; KP.2.9; JN.1; JN.3; JN.1.7; JN.1.32; JN.4; JN.1.41; JN.1.52; JN.1.34; JN.1.17; LV.1; JN.1.4; BA.2.86.1; MC.13; KP.3.2.4; JN.2; JN.16; BA.2.86.2; JQ.1; JN.5; JN.18; JN.3.1; JN.8; JN.7; JN.9; JN.2.1; JN.6 |
| 15 | Omicron: BA.2-like | 02-11-21 | 11-06-25 | Guadeloupe / Basse-Terre | EPI_ISL_13873803 | EPI_ISL_20063868 | BA.2.9; BA.2; BA.2.12.1; BA.2.54; BA.2.44; BA.2.36; BA.2.3; BA.2.52; BA.2.9.3; BA.2.18; BA.2.71; BA.2.2; BA.2.12; BA.2.2.1; BA.2.5; BA.2.10; CB.1; BA.2.9.2; BA.2.10.1; BA.2.10.3; BA.2.37; BA.2.51; BA.2.23; BY.1; BA.2.75.2; BN.1.3; BN.1.3.5; BL.1; BA.2.3.15; BA.2.47 |
| 16 | Omicron: BA.4/BA.5-like | 01-11-21 | 14-06-25 | Slovakia / Prievidza | EPI_ISL_15941105 | EPI_ISL_20070801 | BA.5.2.18; BA.5.5; BA.5.2.20; BA.5.6; BA.5.2.1; BA.4.1; BA.5.1; BA.4.4; BQ.1.1.32; BQ.1.23; BA.5.6.2; BF.7.14; BA.5.3.2; BA.4.1.9; BA.5.1.23; BA.5.2; BQ.1.1; BQ.1; BA.5; BA.5.2.6; BQ.1.1.8; BF.7; BQ.1.1.46; BA.5.2.26; BQ.1.10; BA.4; BA.4.6; BW.1; BA.5.3; CP.1 |
| 17 | Omicron: XBB family | 19-11-21 | 29-04-25 | Slovakia / Okres Zilina | EPI_ISL_18682061 | EPI_ISL_19869479 | XBB.1.16.11; XBB.1.5.99; XBB.2.11; XBB.1.5; XBB.2; XBB; XBB.1.5.79; XBB.1.15; XBB.1; EG.1.3; XBB.1.5.13; FY.6; XBB.1.5.37; XBB.1.5.20; XBB.1.5.2; XBB.1.9.1; FZ.1.1; XBB.1.16; HZ.2; EG.8; XBB.1.43; FE.1.2; EG.13; XBB.1.5.25; FL.1.5.1; XBB.2.11.1; XBB.1.13; XBB.1.16.15; XBB.1.44; XBB.3 |
| 18 | Omicron: recombinant (other) | 03-12-21 | 09-07-25 | Germany / Saxony-Anhalt | EPI_ISL_14463157 | EPI_ISL_20074562 | XAL; XD; XH; XM; XBK; XAR; XJ; XG; XE; XT; XAF; XP; XR; XAC; XAH; XAM; XW; XBF; XAA; XQ; XAN; XAU; XAB; XAZ; XAJ; XAG; XAQ; XAK; XAD; XAT |
| 19 | Omicron: other | 08-11-21 | 06-12-24 | USA / Indiana | EPI_ISL_14847933 | EPI_ISL_19792738 | B.1.1.529; DC.1; CS.1 |
| 20 | Unassigned | 30-12-19 | 21-05-25 | China / Hubei | EPI_ISL_402124 | EPI_ISL_20050307 | B; A; R.1; A.2; B.12; B.33; Unassigned; A.6; B.4; A.19; B.53; B.42; B.43; N.6; B.40; B.5; B.41; B.3; A.16; B.11; B.61; A.1; A.5; B.23; B.31; B.55; B.29; B.6; B.38; B.58 B.1; B.1.1.285; B.1.36.1; B.1.398; B.1.443; B.1.2; B.1.221; B.1.160; B.1.177; B.1.1; B.1.91; B.1.222; B.1.1.331; B.1.356; B.1.1.61; B.1.1.306; B.1.1.323; B.1.1.142; B.1.166; B.1.1.1; B.1.1.28; B.1.1.257; B.1.617; B.1.1.17; B.1.1.204; B.1.147; B.1.36.31; B.1.22; B.1.465; B.1.619 |

**Supplementary Table 3. Annual distribution of spike sequences across countries and regions included in the study.**

| **Country/region** | **2020** | **2021** | **2022** | **2023** | **2024** | **2025** | **Missing/invalid date*** | **Total** |
| --- | --- | --- | --- | --- | --- | --- | --- | --- |
| **Australia** | 371 | 835 | 5,261 | 2,618 | 1,038 | 359 | 38 | 10,520 |
| **Brazil** | 989 | 7,903 | 4,898 | 1,751 | 1,010 | 215 | 170 | 16,936 |
| **Canada** | 1,002 | 6,059 | 8,704 | 7,893 | 5,067 | 822 | 893 | 30,440 |
| **France** | 459 | 10,348 | 11,570 | 5,651 | 2,299 | 357 | 1,132 | 31,816 |
| **Germany** | 529 | 13,861 | 14,015 | 3,930 | 657 | 191 | 108 | 33,291 |
| **India** | 1,186 | 6,129 | 3,398 | 904 | 143 | 42 | 1,914 | 13,716 |
| **Italy** | 463 | 5,126 | 3,774 | 2,359 | 746 | 160 | 310 | 12,938 |
| **Japan** | 930 | 5,097 | 10,651 | 7,121 | 2,245 | 333 | 203 | 26,580 |
| **South Africa** | 410 | 906 | 1,011 | 392 | 173 | 35 | 3 | 2,930 |
| **United Kingdom** | 4,771 | 26,778 | 17,090 | 4,911 | 619 | 106 | 689 | 54,964 |
| **USA California** | 1,348 | 8,213 | 6,225 | 4,346 | 3,014 | 664 | 80 | 23,890 |
| **USA New York** | 472 | 41,00 | 4,485 | 2,970 | 2,116 | 808 | 13 | 14,964 |
| **Total** | **12,930** | **95,355** | **91,082** | **44846** | **19,127** | **4,092** | **5,553** | **272,985** |

* Missing/invalid date = Number of records with invalid dates.

**Supplementary Table 4. Geographic distribution of spike sequences included in the study, stratified by country and region**.

| **Country/region** | **Base dataset** | **Clustering** | **Simulation** | **PCA*** | **Entropy** |
| --- | --- | --- | --- | --- | --- |
| **Australia** | 10,520 | 10,512 | 10,474 | 10,201 | 10,159 |
| **Brazil** | 16,936 | 16,756 | 16,589 | 16,080 | 15,876 |
| **Canada** | 30,440 | 30,052 | 29,185 | 29,655 | 28,792 |
| **France** | 31,816 | 31,690 | 30,741 | 31,260 | 30,281 |
| **Germany** | 33,291 | 33,220 | 33,112 | 32,687 | 32,584 |
| **India** | 13,716 | 13,689 | 11,776 | 13,201 | 11,417 |
| **Italy** | 12,938 | 12,880 | 12,570 | 12,630 | 12,302 |
| **Japan** | 26,580 | 26,465 | 26,265 | 26,019 | 25,814 |
| **South Africa** | 2,930 | 2,906 | 2,903 | 2,795 | 2,792 |
| **United Kingdom** | 54,964 | 54,882 | 54,193 | 54,299 | 53,649 |
| **USA California** | 23,890 | 23,830 | 23,750 | 23,290 | 23,205 |
| **USA New York** | 14,964 | 14,887 | 14,873 | 14,216 | 14,203 |
| **TOTAL** | **272,985** | **271,769** | **266,431** | **266,333** | **261,074** |

* PCA, principal component analysis.

**Supplementary Table 5. Pairwise multivariate analysis of variance (MANOVA) comparisons between physicochemical clusters, with *p* values adjusted for multiple testing using the Benjamini–Hochberg procedure.**

| **Cluster 1** | **Cluster 2** | **N1** | **N2** | **Test** | **Statistic** | **Approximate F** | **Df1** | **Df2** | **BH-adjusted p** |
| --- | --- | --- | --- | --- | --- | --- | --- | --- | --- |
| 1 | 2 | 150211 | 66175 | Pillai | 0.685385769 | 157129.0903 | 3 | 216382 | <10^-6^ |
| 1 | 3 | 150211 | 33919 | Pillai | 0.861603106 | 382098.0096 | 3 | 184126 | <10^-6^ |
| 1 | 4 | 150211 | 5878 | Pillai | 0.278583912 | 20091.39647 | 3 | 156085 | <10^-6^ |
| 1 | 5 | 150211 | 5863 | Pillai | 0.333498347 | 26030.98675 | 3 | 156070 | <10^-6^ |
| 1 | 6 | 150211 | 4287 | Pillai | 0.507805134 | 53131.29127 | 3 | 154494 | <10^-6^ |
| 2 | 3 | 66175 | 33919 | Pillai | 0.932356926 | 459862.8227 | 3 | 100090 | <10^-6^ |
| 2 | 4 | 66175 | 5878 | Pillai | 0.88452405 | 183960.5943 | 3 | 72049 | <10^-6^ |
| 2 | 5 | 66175 | 5863 | Pillai | 0.686060161 | 52472.53495 | 3 | 72034 | <10^-6^ |
| 2 | 6 | 66175 | 4287 | Pillai | 0.880508077 | 173062.8493 | 3 | 70458 | <10^-6^ |
| 3 | 4 | 33919 | 5878 | Pillai | 0.956099901 | 288883.8087 | 3 | 39793 | <10^-6^ |
| 3 | 5 | 33919 | 5863 | Pillai | 0.664623525 | 26276.33573 | 3 | 39778 | <10^-6^ |
| 3 | 6 | 33919 | 4287 | Pillai | 0.768177934 | 42196.05997 | 3 | 38202 | <10^-6^ |
| 4 | 5 | 5878 | 5863 | Pillai | 0.955145635 | 83310.68925 | 3 | 11737 | <10^-6^ |
| 4 | 6 | 5878 | 4287 | Pillai | 0.968394799 | 103778.9056 | 3 | 10161 | <10^-6^ |
| 5 | 6 | 5863 | 4287 | Pillai | 0.873005711 | 23249.11873 | 3 | 10146 | <10^-6^ |

N1 and N2 denote the numbers of spike sequences assigned to the first and second clusters in each pairwise compariosn, respectively. The test statistic is Pillai’s trace and the approximate F statistic is reported. Df1 and Df2 denote the numerator and dinominator degrees of freedom of the approximate F statistic, respectively. Adjusted p-values were calculated using the Benjamini–Hochberg procedure.

**Supplementary Table 6. Bootstrap estimates of cluster centroid stability for the three standarized physicochemical properties.** Clustter centroids were recalculated across 200 bootstrap resamples. Mean centroid estimates, bootstrap standard deviations (SD), and 95% percentile confidence intervals (2.5^th^ -97.5^th^ percentiles) are reported for each cluster.

|  | **Charge (pH=7.0)** | | | |
| --- | --- | --- | --- | --- |
| **Cluster** | **Mean** | **SD** | **CI (0.025)** | **CI (0.975)** |
| **1** | 0.583 | 0.001 | 0.581 | 0.585 |
| **2** | -0.190 | 0.001 | -0.193 | -0.188 |
| **3** | -2.022 | 0.001 | -2.025 | -2.019 |
| **4** | 1.151 | 0.003 | 1.144 | 1.157 |
| **5** | -0.990 | 0.003 | -0.995 | -0.983 |
| **6** | -1.741 | 0.004 | -1.749 | -1.733 |
|  | **Hydropathy** | | | |
| **Cluster** | **Mean** | **SD** | **CI (0.025)** | **CI (0.975)** |
| **1** | 0.213 | 0.002 | 0.210 | 0.217 |
| **2** | -1.139 | 0.002 | -1.144 | -1.135 |
| **3** | 0.697 | 0.003 | 0.691 | 0.702 |
| **4** | 0.360 | 0.006 | 0.348 | 0.372 |
| **5** | 0.643 | 0.009 | 0.626 | 0.663 |
| **6** | 2.099 | 0.007 | 2.085 | 2.112 |
|  | **Aliphatic index** | | | |
| **Cluster** | **Mean** | **SD** | **CI (0.025)** | **CI (0.975)** |
| **1** | -0.046 | 0.002 | -0.050 | -0.041 |
| **2** | 0.148 | 0.003 | 0.142 | 0.154 |
| **3** | 0.388 | 0.004 | 0.379 | 0.396 |
| **4** | -2.475 | 0.007 | -2.490 | -2.461 |
| **5** | 0.046 | 0.012 | 0.023 | 0.069 |
| **6** | -0.695 | 0.008 | -0.711 | -0.678 |

**Supplementary Table 7. Cross-validated confusion matrix for linear discriminant analysis (LDA) classification based on physicochemical variables.** Cluster 0 (Ν = 5,436) was excluded from the analysis. The overall cross-validated classification accuracy exceeded 96%.

| **Cluster observed** | **Cluster predicted** | **N** |
| --- | --- | --- |
| 1 | 1 | 141,597 |
| 2 | 2 | 66,174 |
| 3 | 3 | 33,919 |
| 4 | 4 | 5,878 |
| 1 | 2 | 5,838 |
| 5 | 5 | 4,836 |
| 6 | 6 | 4,173 |
| 1 | 4 | 1,942 |
| 1 | 5 | 834 |
| 5 | 3 | 507 |
| 5 | 2 | 480 |
| 6 | 3 | 114 |
| 5 | 6 | 40 |
| 2 | 1 | 1 |
| **Total** |  | **266,333** |

**Supplementary Table 8. Bootstrapped means and standard deviations (SDs) of physicochemical properties (charge at pH 7.0, hydropathy, and aliphatic index) for each cluster.** Values are reported as mean ± SD, with the corresponding minimum and maximum values provided in parentheses.

| **Physiochemical property** | | | |
| --- | --- | --- | --- |
| **Cluster** | **Charge at pH=7 (min, max)** | **Hydropathy (min, max)** | **Aliphatic index (min, max)** |
| **1** | 2.83 ± 0.93 (-0.10, 6.13) | -0.077 ± 0.004 (-0.092, -0.062) | 84.69 ± 0.30 (83.63, 85.73) |
| **2** | 1.14 ± 0.624 (-1.11,3.13) | -0.085± 0.004 (-0.098, -0.073) | 84.76 ± 0.27 (83.81, 85,67) |
| **3** | -2.71 ± 0.72 (-4.63, -1.35) | -0.074 ± 0.003 (-0.086, -0.063) | 84.84 ± 0.27 (83.95, 85.74) |
| **4** | 4.06 ± 0.57 (2.65, 5.37) | -0.076 ± 0.003 (-0.085, -0.068) | 83.90 ± 0.19 (83.37, 84.34) |
| **5** | -0.59 ± 0.51 (-1.84, 0.37) | -0.075 ± 0.004 (-0.086, -0.063) | 84.72 ± 0.29 (83.98, 85.44) |
| **6** | -2.22 ± 0.60 (-3.87, -0.59) | -0.066 ± 0.003 (-0.074, -0.059) | 84.48 ± 0.17 (83.98. 84.98) |

**Supplementary Table 9. Summary statistics of monthly genotype–phenotype entropy decoupling across the 12 geographic regions.** The entropy difference (ΔH) was calculated as normalized phenotypic Shannon entropy minus normalized genotypic Shannon entropy. Mean ΔH indicates the overall direction of the difference, whereas mean, median, maximum and 95^th^ percentile of absolute differences (|ΔH|) quantify its magnitude irrespective of direction. The percentage of months with ΔH < 0 indicates the frequency of months in which normalized genotypic entropy exceeded normalized phenotypic entropy. Regions are ranked in descending order of mean absolute entropy difference (mean |ΔH|), reflecting the average magnitude of genotype-phenotype entropy decoupling.

| **Rank** | **Country/Region** | **Months** | **Mean ΔH** | **Mean \|ΔH\|** | **Median \|ΔH\|** | **Max \|ΔH\|** | **95^th^ percentile \|ΔH\|** | **ΔH < 0 (%)** |
| --- | --- | --- | --- | --- | --- | --- | --- | --- |
| 1 | Australia | 53 | -0.193 | 0.411 | 0.378 | 0.93 | 0.888 | 64.2 |
| 2 | USA (California) | 64 | -0.052 | 0.351 | 0.345 | 0.901 | 0.746 | 50 |
| 3 | Canada | 62 | -0.093 | 0.335 | 0.325 | 0.883 | 0.794 | 50 |
| 4 | Germany | 53 | -0.081 | 0.333 | 0.253 | 0.983 | 0.791 | 58.5 |
| 5 | France | 61 | -0.094 | 0.332 | 0.275 | 0.882 | 0.815 | 49.2 |
| 6 | Italy | 51 | -0.088 | 0.315 | 0.265 | 0.875 | 0.834 | 54.9 |
| 7 | United Kingdom | 56 | -0.058 | 0.31 | 0.321 | 0.788 | 0.695 | 42.9 |
| 8 | USA (New York) | 59 | -0.007 | 0.301 | 0.296 | 0.815 | 0.641 | 49.2 |
| 9 | Japan | 59 | -0.149 | 0.287 | 0.229 | 0.891 | 0.817 | 52.5 |
| 10 | India | 42 | 0.068 | 0.285 | 0.228 | 0.974 | 0.757 | 31 |
| 11 | Brazil | 58 | 0.093 | 0.28 | 0.236 | 0.789 | 0.649 | 34.5 |
| 12 | South Africa | 32 | -0.053 | 0.228 | 0.2 | 0.604 | 0.501 | 53.1 |
